# The roles of an extended N-terminal region and ETD motif in a pump-like cation channelrhodopsin discovered in a lake microbiome

**DOI:** 10.1101/2024.05.16.594411

**Authors:** Shunki Takaramoto, Shai Fainsod, Takashi Nagata, Andrey Rozenberg, Oded Béjà, Keiichi Inoue

## Abstract

Channelrhodopsins are light-gated ion channels consisting of seven-transmembrane helices and a retinal chromophore, which are used as popular optogenetic tools for modulating neuronal activity. Cation channelrhodopsins (CCRs), first recognized as the photoreceptors in the chlorophyte *Chlamydomonas reinhardtii*, have since been identified in diverse species of green algae, as well in other unicellular eukaryotes. The CCRs from non-chlorophyte species are commonly referred to as bacteriorhodopsin-like channelrhodopsins, or BCCRs, as most of them feature the three characteristic amino acid residues of the “DTD motif” in the third transmembrane helix (TM3 or helix C) matching the canonical DTD motif of the well-studied archaeal light-driven proton pump bacteriorhodopsin. Here, we report characterization of HulaCCR1, a novel BCCR identified through metatranscriptomic analysis of a unicellular eukaryotic community in Lake Hula, Israel. Interestingly, HulaCCR1 has an ETD motif in which the first residue of the canonical motif is substituted for glutamate. Electrophysiological measurements of the wild-type and a mutant with a DTD motif of HulaCCR1 suggest the critical role of the first glutamate in spectral tuning and channel gating. Additionally, HulaCCR1 exhibits long extensions at the N– and C-termini. Photocurrents recorded from a truncated variant without the signal peptide predicted at the N-terminus were diminished, and membrane localization of the truncated variant significantly decreased, indicating that the signal peptide is important for membrane trafficking of HulaCCR1. These characteristics of HulaCCR1 would be related to a new biological significance in the original unidentified species, distinct from those known for other BCCRs.

## Introduction

Microbial rhodopsins are a family of photoreactive membrane proteins that have seven transmembrane helices and a retinal chromophore covalently binding to a conserved lysine residue in the 7th transmembrane helix (called TM7 or helix G) [1–3]. Among them, channelrhodopsins (ChRs) are light-induced ion channels and are extensively utilized in optogenetics [4, 5]. Channelrhodopsins were first discovered in green algae when the photoreceptors responsible for phototactic behavior in *Chlamydomonas reinhardtii* were identified as microbial rhodopsin channels transporting cations, or cation channelrhodopsins (CCRs) [6–8]. In contrast to *Chlamydomonas*, two different types of ChRs were subsequently identified in the cryptophyte *Guillardia theta*: a distinct family of CCRs, as well as a family of anion-transporting ChRs, or anion channelrhodopsins (ACRs), which was the first indication of the high diversity of ChRs among unicellular eukaryotes [9, 10]. Natural and engineered CCRs and ACRs have since been used as molecular tools in optogenetic research to stimulate or inhibit neuronal firing by light, respectively [11–14].

Many microbial rhodopsins, including the well-characterized light-driven proton pump bacteriorhodopsin from *Halobacterium salinarum* (*<u>Hs</u>*BR), feature three characteristic amino acid residues together constituting the canonical “DTD motif” in the third transmembrane helix (called TM3 or helix C). This motif is composed of two aspartic acid residues and a threonine residue: D85/D96 and T89 in *Hs*BR, respectively [15]. One remarkable feature of the cryptophyte CCRs from *Guillardia theta* was the presence of the DTD motif identical to that of *Hs*BR [10, 16]. Following the discovery of the CCRs from *G. theta*, many CCRs possessing the DTD or similar motifs were found in other species, collectively referred to as the bacteriorhodopsin-like channelrhodopsins (BCCRs). Among them, ChRmine from the cryptophyte *Rhodomonas lens* exhibiting high photocurrents and high light sensitivity has emerged as the model BCCR and has found its application in deep transcranial optogenetics [17–19]. Recent years have also witnessed the discovery of highly potassium selective BCCRs, the kalium channelrhodopsins (KCRs) from the stramenopiles *Hyphochytrium catenoides* (*Hc*KCR1 and *Hc*KCR2) [20] and *Wobblia lunata* (WiChR) [21], as well as other heterotrophic flagellates. Since KCRs efficiently induce hyperpolarization by K^+^ efflux, they are used for precise optogenetic inhibition.

Here, we characterize HulaCCR1, a novel microbial rhodopsin identified through metatranscriptomic analysis of the unicellular eukaryotic community of Lake Hula, Israel. This rhodopsin has an ETD motif in TM3 and belongs to the family of stramenopile and other heterotrophic flagellate CCRs (SoHF CCRs) which includes KCRs and other CCRs from cultured unicellular eukaryotes and environmental sources [21, 22] (Figs. 1a and b). While uncommon among BCCRs, the ETD motif is not an idiosyncrasy of HulaCCR1 and is found in the related proteins mgCCR1 [22] and P1ChR1 [21]. The structure of HulaCCR1 predicted with AlphaFold2 Colab [23] features an unusually long N-terminal extension with a signal peptide and a prominent partially structured long C-terminal extension (Fig. 1c).

**Fig. 1.**
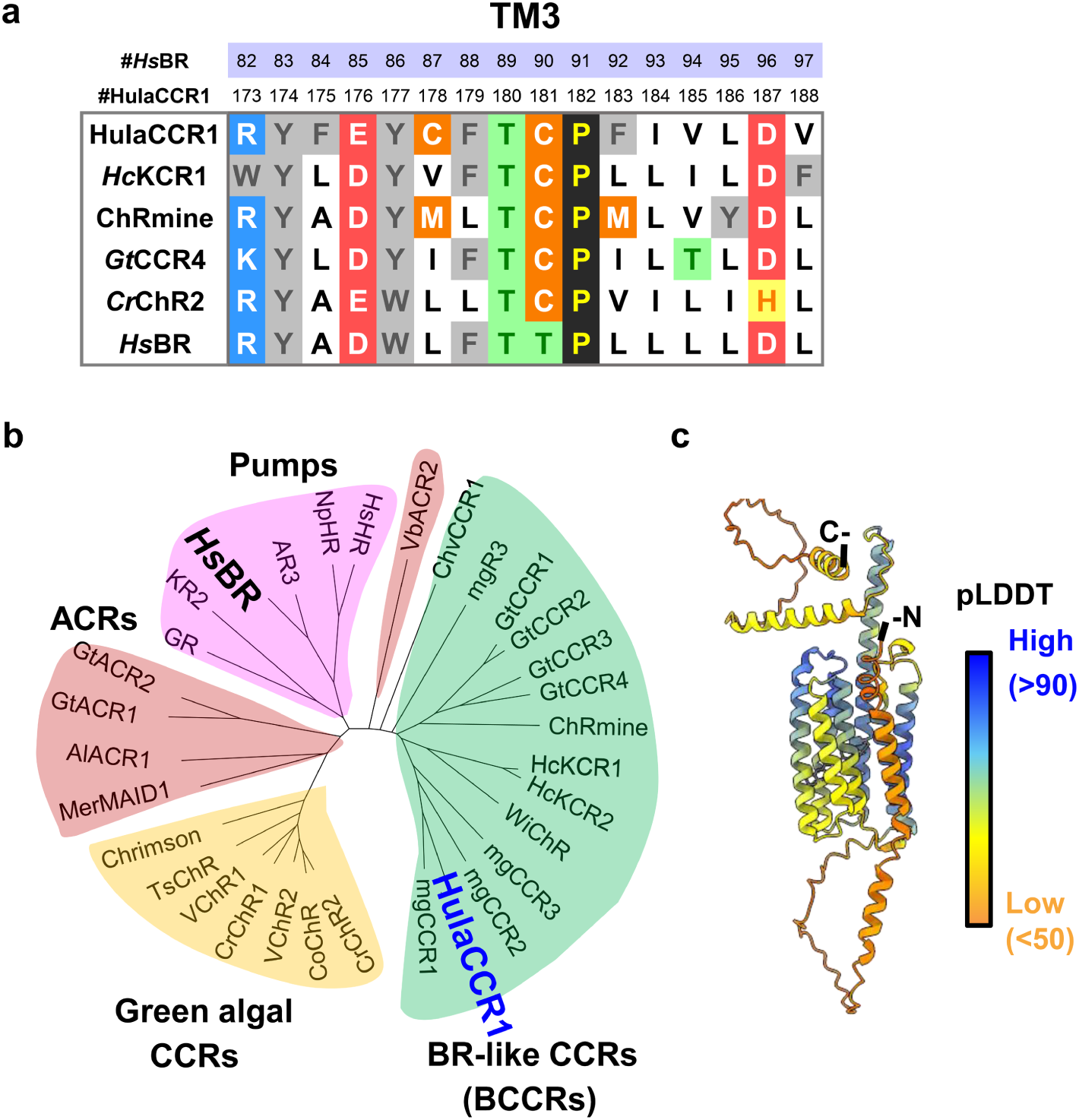
Amino acid sequence alignment of HulaCCR1, phylogenetic relationship, and predicted structure. **a** Amino acid alignment of TM3 of HulaCCR1 with *Hc*KCR1, ChRmine, *Gt*CCR4, *Cr*ChR2, and *Hs*BR. **b** The phylogenetic tree of representative channelrhodopsins, ion-pumping rhodopsins, and HulaCCR1. The phylogenetic tree was inferred using the neighbor-joining method of MEGA X software (version 11.0.13). **c** Schematic illustration of HulaCCR1 structure based on AlphaFold2 prediction.

In this study, we investigated the function of HulaCCR1 using electrophysiological techniques. As a result, it was revealed that this rhodopsin exhibits non-specific cation channel activity. To understand the role of the substitution of glutamate for the ancestral aspartate at the first position of the TM3 motif in HulaCCR1, we examined the effects of this residue on the channel activity. In addition, we investigated the role of the long extensions at the N– and C-termini of HulaCCR1. Analyses of the distribution and origins of the clade of ChRs to which HulaCCR1 belongs, show that it can be subdivided into six subclades with the freshwater subclades found in lakes on both sides of the Atlantic and originating from the heterotrophic flagellate group of kathablepharids.

## Results

### Identification of HulaCCR1

HulaCCR1 was identified as part of an ongoing metatranscriptomic study of the unicellular eukaryotic community of Lake Hula in Northern Israel. This protein is representative of a distinct clade within the family of stramenopile and other heterotrophic flagellate CCRs (SoHF CCRs) to which the partially characterized ChRs mgCCR1–3 [22] also belong (Fig. 1b, Figs. 2a and b). Proteins from this clade were found in a number of other metatranscriptomic and metagenomic datasets and their diversity could be circumscribed to four subclades appearing in freshwater datasets and two subclades in marine datasets: freshwater subclades A to C include mgCCR1, HulaCCR1 and mgCCR2, respectively and marine subclade A includes mgCCR3 (see Fig. 2b). Freshwater subclades A and B are the closest relatives, with a modest overall pairwise sequence identity of 48% between HulaCCR1 and mgCCR1, least similar in the N-terminal extension. Ideal for the study of the role of the carboxylate amino acid at the first position of the TM3 motif, this clade demonstrates both DTD and ETD variants, with the mgCCR1 and HulaCCR1 subclades having the otherwise uncommon glutamate.

**Fig. 2.**
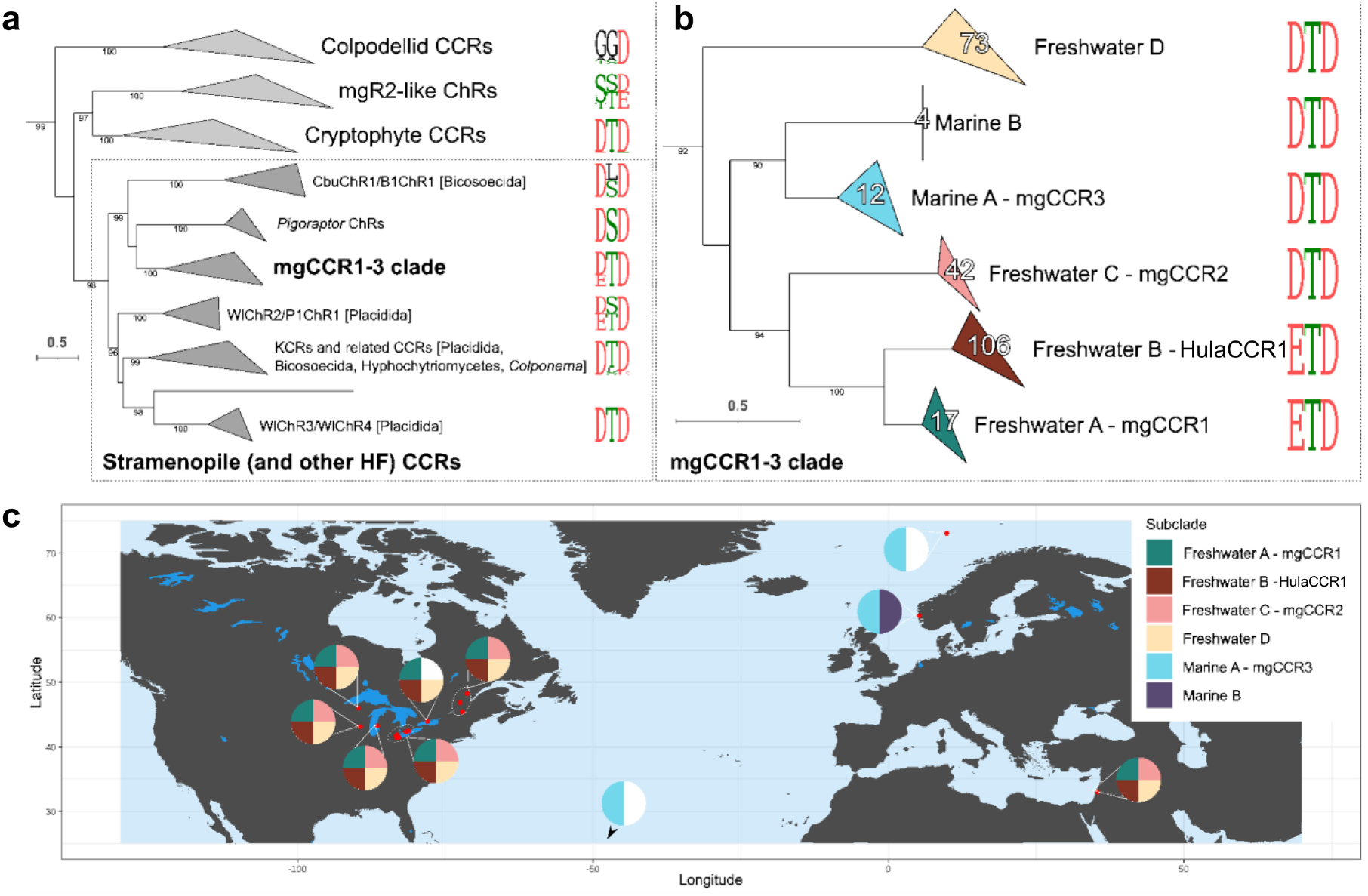
Phylogenetic relationships of HulaCCR1 and the mgCCR1-3 clade and their geographical distribution. **a** Position of the mgCCR1-3 clade among the stramenopile (and other heterotrophic flagellate) CCRs and BCCRs. The BCCR superclade was extracted from the ChR phylogeny in [28], numbers below the branches indicate ultra-fast bootstrap support values ≥95. **b** Fine phylogenetic relationships among the six subclades of the mgCCR1-3 clade. The tree is outgroup-rooted with the outgroups not shown. Numbers below the branches connecting subclades indicate rapid bootstrap support values ≥50. Numbers inside the triangles correspond to the number of the metagenomic sequences placed on the tree (see Materials and Methods for details). Weblogos of the TM3 motif are provided for the BCCR clades and subclades in **a** and **b**. **c** Distribution of the four freshwater and two marine subclades of the mgCCR1-3 clade based on environmental assemblies in JGI/IMG and Lake Hula. The pie charts show presence/absence of the corresponding four (freshwater locations) or two (marine locations) subclades. For the list of the datasets see Fig. S2.

Since none of the members of the mgCCR1-3 clade are known from cultured creatures and the available genomic fragments from the metagenomic datasets provide only a meager genomic background for the ChR genes, we attempted to identify their origin by focusing on potential mobile elements which might be shared with sequenced genomes. To achieve this, we took DNA sequences immediately upstream and downstream of the ChR genes and recruited other genomic fragments with high nucleotide similarity from the same environmental samples. This strategy indeed yielded multiple genomic fragments containing retrotransposons, all of which belonged to the Ty3/gypsy group. While one of the contigs containing a ChR gene from the freshwater subclade D itself contained a fragment of such retrotransposon, related retrotransposons were also found in contigs recruited with regions flanking subclade-A and C genes as well, which indicates that they might overlap retrotransposon integration sites. Phylogenetic analysis showed all of these elements to belong to the same pool of retrotransposons shared with Ty3 elements from four marine kathablepharid single amplified genomes (SAGs) [24] (Fig. S1a). This strongly indicates that at least the freshwater subclades originate from kathablepharids, a group of cryptist heterotrophic flagellates related to cryptomonads [25, 26]. Analysis of the small subunit rRNA (SSU) gene fragments revealed that kathablepharids are widely attested in the samples in which the mgCCR1-3 clade is found and are one of the most typical components of the eukaryotic communities in them (Fig. S2). Phylogenetic analysis of a representative complete kathablepharid SSU type dominant in these samples, places it in the freshwater kathablepharid clade [27] (Fig. S1b). While among cryptist groups, ChRs are known from the photosynthetic cryptophyceans (cryptophyte ACRs [9] and CCRs [10]) as well as from the heterotrophic *Hemiarma marina* (same two families) and *Goniomonas pacifica* (an mgR2-like ChR and the unique *Gp*ACR1 [28]), the very limited genetic data available for kathablepharids have thus revealed only a putative mgR2-like CCR in the transcriptome of *Roombia truncata* [29] (Fig. S1b). The kathablepharid origin of the mgCCR1-3 clade significantly expands both the repertoire of the ChRs among cryptists and the distribution of the SoHF CCRs.

### Ion transport activity of HulaCCR1

Upon establishing the expression of HulaCCR1 in COS-1 cells, we investigated the absorption spectra of the protein solubilized from the membrane fraction of the cells with a solution containing *n*-dodecyl-β-D-maltoside (DDM). The absorption maximum wavelength (*λ*_max_) of HulaCCR1 was determined to be 545 nm by a hydroxylamine (HA) bleaching method [30] (Fig. 3a). To investigate the molecular function of HulaCCR1, electrophysiological measurements were conducted. Based on the strong EYFP fluorescence in the outlines of transfected cultured cells, we determined that HulaCCR1 was well expressed in the plasma membrane (Fig. 3b). Whole-cell patch clamp recordings were performed using these cells. With standard pipette ([K^+^] = 128 mM) and extracellular ([Na^+^] = 142 mM) solutions, a significant photocurrent appeared upon illumination with 542-nm light (Fig. 3c, left). The photocurrent depended on the holding potential and exhibited an inversion from positive to negative currents along with increasing the holding potential from −80 to 60 mV, indicating that HulaCCR1 exhibits light-dependent ion channel activity. Changing the cations in both the patch pipette and extracellular solutions to N-methyl-D-glucamine (NMG) resulted in a profound decrease in photocurrent intensity (Fig. 3c, right). Furthermore, in this measurement, despite the anion conditions being asymmetric between the external and internal side of the membrane, where the pipette and extracellular solutions contained 90 mM glutamate and 151 mM Cl^−^, respectively, the reversal potential remained at around 0 mV, excluding the possibility of anion channel activity and indicating H^+^ channel activity. We conclude that HulaCCR1 possesses cation channel transport activity.

**Fig. 3.**
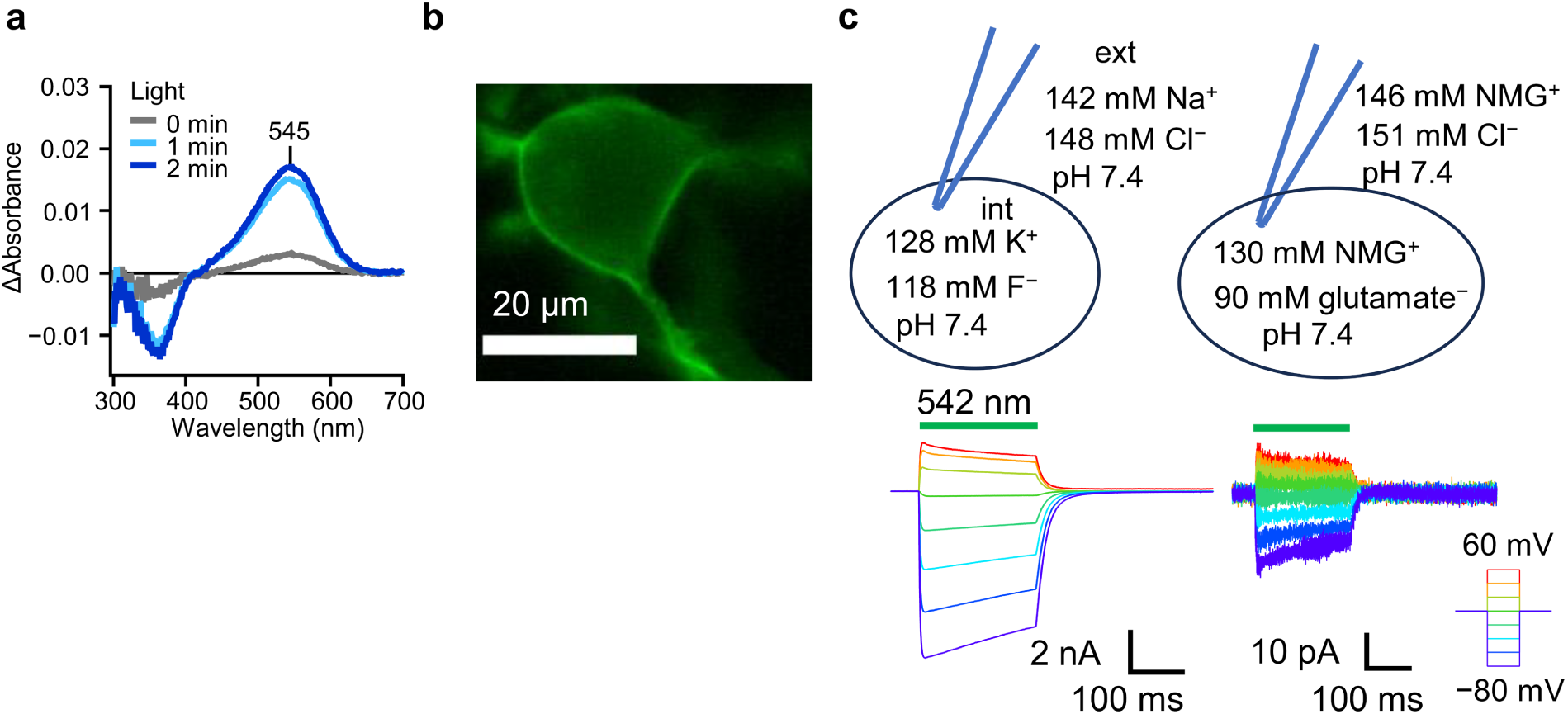
Characterization of HulaCCR1. **a** Difference UV–visible absorption spectra between before and after the hydroxylamine bleach of the HulaCCR1 showing the *λ*_max_ at 545 nm. The difference spectra were measured upon the hydrolysis reaction of the retinal Schiff-base linkage induced by hydroxylamine with visible light illumination (0, 1, or 2 min). **b** Fluorescence image of ND7/23 cell expressing HulaCCR1 which is C-terminally labeled with EYFP (Scale bar: 20 μm). **c** Photocurrents of HulaCCR1 at 142 mM [Na^+^]_out_ and 128 mM [K^+^]_in_ at pH = 7.4 (left) and at 146 mM [NMG^+^]_out_ and 130 mM [NMG^+^]_in_ at pH = 7.4 (right).

Next, to investigate the ion selectivity of HulaCCR1, photocurrents were measured in extracellular solutions containing different cations (Fig. 4a), and current–voltage (*I*–*V*) curves were plotted based on peak current intensities (Fig. 4b). The *I*–*V* curves were fitted by a quadratic function, then the reversal potentials (*E*_rev_) of the photocurrents were estimated. Taking the results with NMG-containing extracellular solution as a reference, the reversal potential shift (Δ*E*_rev_) was calculated. Δ*E*_rev_ values increased with monovalent cations, while only low values were observed with divalent cations, Mg^2+^ and Ca^2+^. Hence, Mg^2+^ and Ca^2+^ hardly permeate through the channel pore in HulaCCR1. Assuming that only monovalent cations in the extracellular/pipette solutions permeate, the relative cation permeability (*P*_X+_/*P*_H+_) was calculated using Goldman–Hodgkin–Katz (GHK) equation (eq. 1) from Δ*E*_rev_ values,

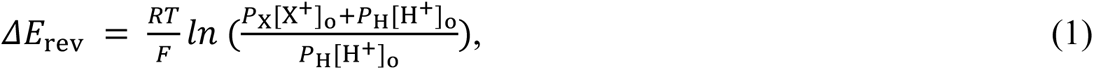

**Fig. 4.**
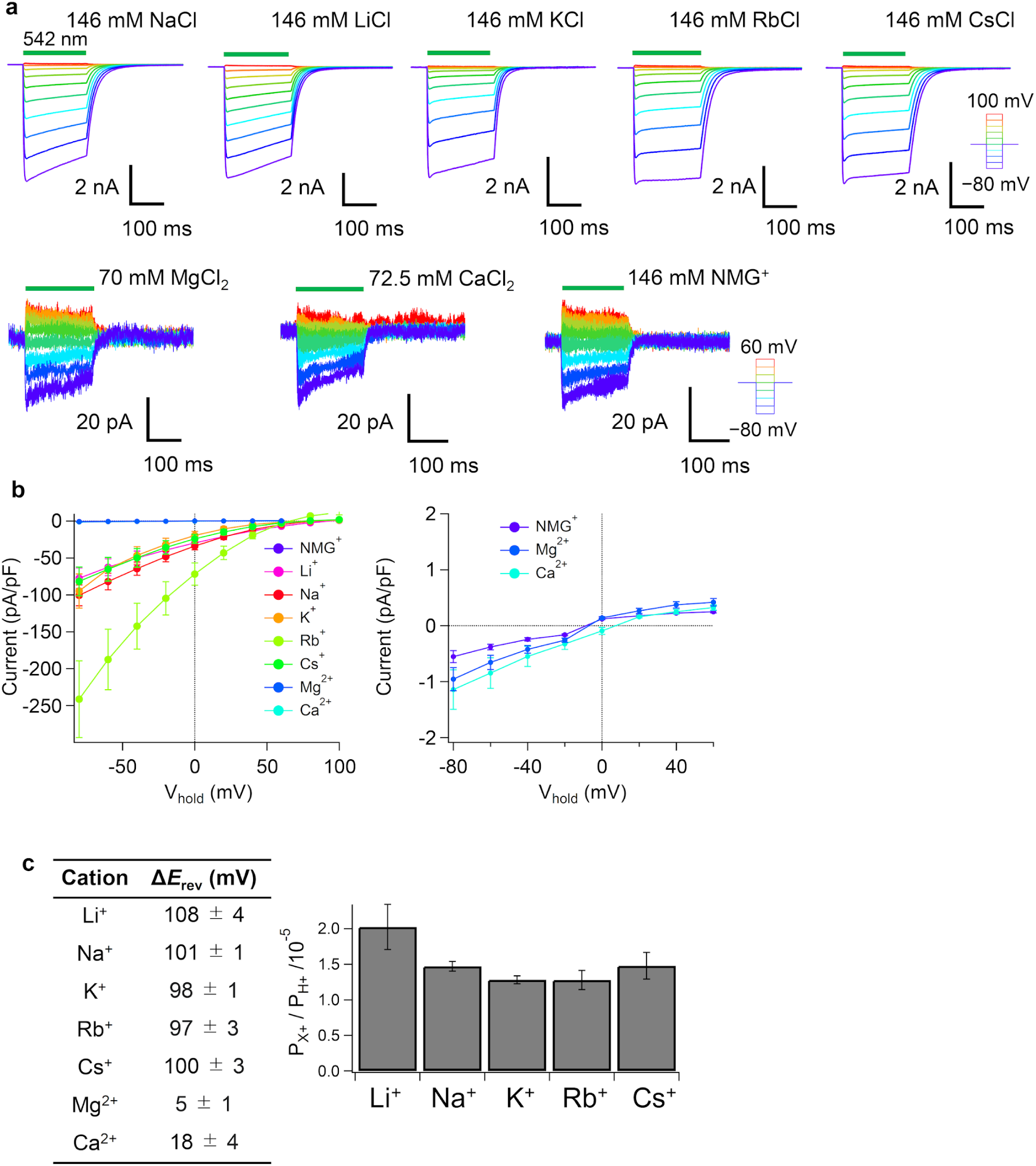
Ion selectivity of HulaCCR1. **a** Representative photocurrent traces of HulaCCR1 at 146 mM [X^+^]_out_ and 130 mM [NMG^+^]_in_ (top), and at 70 mM [Mg^2+^]_out_ /72.5 mM [Ca^2+^]_out_ /146 mM [NMG^+^]_out_ and 130 mM [NMG^+^]_in_ (bottom). **b** The *I*–*V* plot of HulaCCR1 photocurrents with different extracellular cations (mean ± S.E., *n* = 6). **c** Difference in *E*_rev_ between different cations and H^+^ (Δ*E*_rev_) (left) and relative permeability of different cations against H^+^ (*P*_X+_/*P*_H+_) (right) (mean ± S.E., *n* = 6).

where *R* is the gas constant, *T* is the temperature, and *F* is the Faraday constant. The obtained values *P*_X+_/*P*_H+_ are shown in Fig. 4c. *P*_Mg2+_/*P*_H+_ and *P*_Ca2+_/*P*_H+_ were not determined by GHK flux equation [31] for divalent cations at equilibrium (eq. 2) because the calculated values of *P*_Mg2+_/*P*_H+_ and *P*_Ca2+_/*P*_H+_ were considerably smaller than the experimental error.

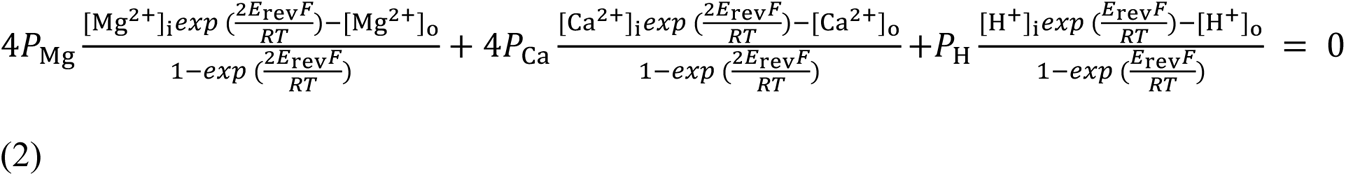

From these results, it can be concluded that HulaCCR1 exhibits a strong ion selectivity for monovalent cations, and this property is similar to those of *Gt*CCR4 [32] and ChRmine [33]. Moreover, there was not much difference in the selectivity among monovalent alkali metal cations (Fig. 4c).

### Mutational analysis of spectral tuning and gating kinetics in HulaCCR1

Many BCCRs share the DTD motif in TM3 identical to the residues D85, T89, and D96 of *Hs*BR. As mentioned above, in HulaCCR1, a glutamate (E176) is present at the position of the first motif residue, forming an ETD motif (Figs. 1a, 2b and S3). To investigate the roles of the amino acid residues near the retinal chromophore, including E176 (Fig. 5a), photocurrents of several single mutants were measured (Figs. 5b–e and S4a). While the E176D mutant, in which the DTD motif is restored, exhibited photocurrents similar to those of the wild-type protein (WT), its action spectrum slightly blue-shifted compared to the WT spectrum (Fig. 5b), indicating the red-shifting spectral-tuning effect of E176 in HulaCCR1. To determine whether this residue is crucial for channel gating, photocurrents were measured with a higher time resolution using nanosecond-pulsed laser flashes (Fig. 5c). The time course of photocurrents due to channel opening and closing was reproduced with a quadruple-exponential function. The obtained time constants were shown in Fig. 5d. Notably, the closing rate of outward current in E176D was significantly slower compared to that of the WT, suggesting the importance of E176 in the closing kinetics.

**Fig. 5.**
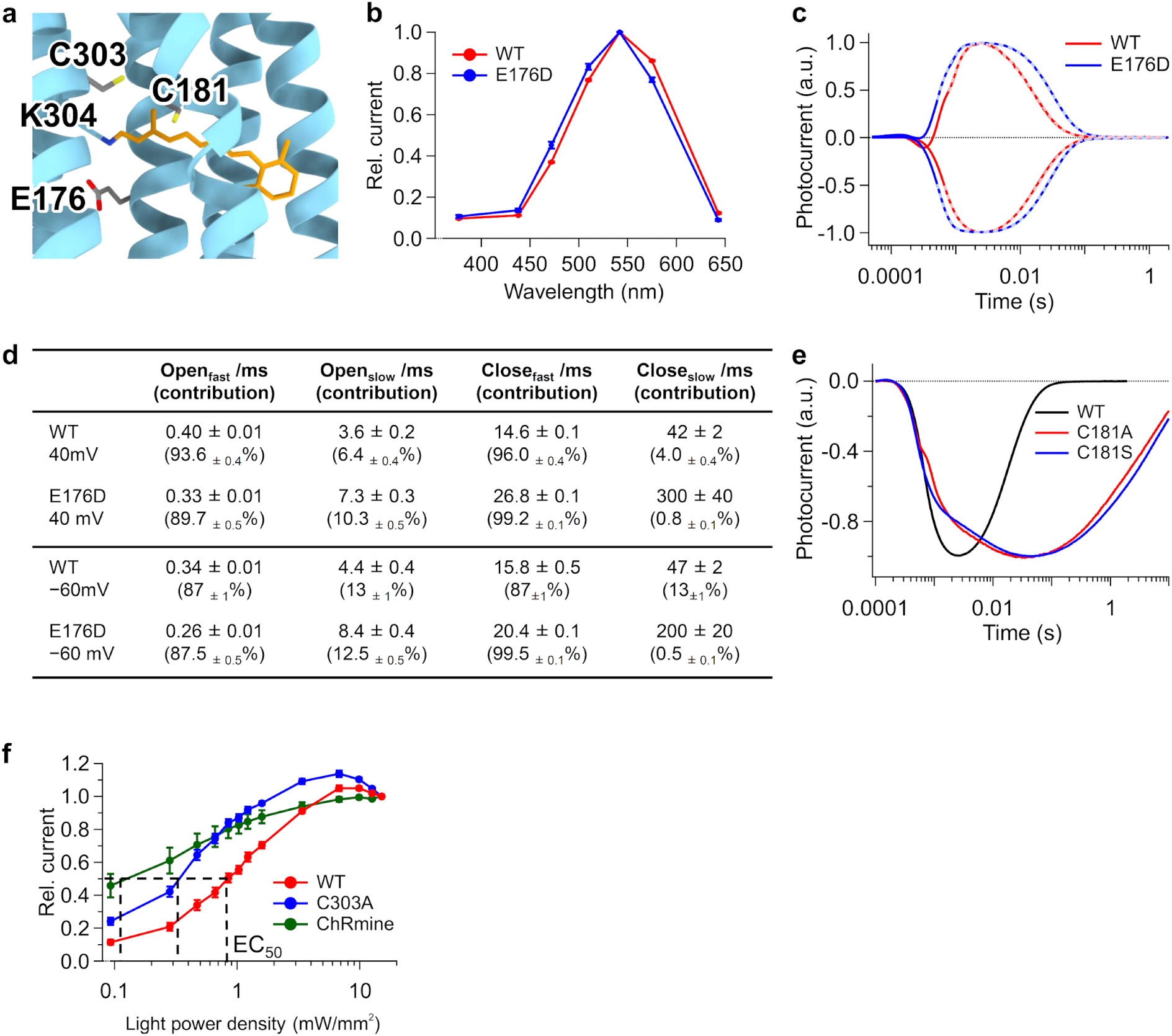
Effect of point mutations. **a** AlphaFold2 predicted structural model for HulaCCR1 showing the mutated residues. The retinal structure (orange) and the side chain of conjugated lysine are of *Hc*KCR1 (PDB ID: 8H86). **b** Normalized action spectra of the HulaCCR1 WT and E176D (mean ± S.E., *n* = 5). **c** Photocurrent traces of the WT and E176D upon nanosecond laser flash excitation at holding potentials of 40 mV (positive) and −60 mV (negative). Photocurrents were recorded at 133 mM [Na^+^]_out_ and 120 mM [Na^+^]_in_ at pH = 7.4. Fitting curves are shown by broken lines. **d** Time constants of channel opening and closing of the WT HulaCCR1 and E176D (mean ± S.E., *n* = 9). **e** Photocurrent traces of the HulaCCR1 WT, C181A, and C181S upon nanosecond laser flash excitation recorded with standard pipette and extracellular solutions. **f** Light-power dependency of the photocurrent of HulaCCR1 WT, C303A, and ChRmine at −40 mV with standard pipette and extracellular solutions (mean ± S.E., *n* = 5).

In chlorophyte channelrhodopsins and BCCRs, a cysteine residue is conserved at the position corresponding to T90 in *Hs*BR in TM3 (Fig. 1a). This cysteine together with D156 in TM4 constitutes the “DC gate” in *Cr*ChR2 [34], and HulaCCR1 mutants in which C181 were substituted with alanine or serine (C181A and C181S) exhibited significantly slower channel closing after 200-ms illumination (Fig. S4a). Both mutants showed a highly prolonged channel closing rate in the nanosecond-pulse excitation experiments (Fig. 5e), as observed in cysteine mutants of other channelrhodopsins [35, 36].

Next, we attempted to tune the spectrum of HulaCCR1 by mutating amino acid residues near the retinal chromophore. First we noticed that HulaCCR1 has C303 adjacent to K304 that forms the Schiff-base linkage with the retinal chromophore in TM7 (Fig. 5a), a trait shared most notably with the KCRs (Fig. S3) [20], while most of the other BCCRs have alanine at this position similarly to *Hs*BR. Importantly, in the case of *Cr*ChR2, substituting alanine for serine at this position causes a red shift in the absorption spectrum [37]. Nevertheless, the action spectrum of the photocurrents of the HulaCCR1 C303A mutant showed no significant red shift compared to the WT (Fig. S4c). Taking a different approach, we attempted to introduce negative charges on the β-ionone ring side of the retinal which is known to stabilize the electronically excited state, thus leading to a red shift in the absorption spectrum [38]. Hence, the action spectra of mutants of residues shown in Fig. S4b, L208M, A232C, A232S, and P276T were measured. This strategy did result in significantly red-shifted action spectra compared to that of the WT (Fig. S4c).

Although the expected spectral shift did not occur in C303A, we found a significant difference in other photocurrent behaviors in this mutant. The decay rate of photocurrents after light-off in C303A was slower compared to that of the WT (Figs. S4a and d). The slower channel closing might be the result of a slower photocycle, as seen for example in DC-gate mutants of *Cr*ChR2 [35, 39]. The dependence of photocurrent intensity on excitation light power was also investigated under continuous light illumination. In this case, it is anticipated that for channelrhodopsins with slower photoreaction cycles, the photocurrent intensity will saturate at a lower light intensity compared to those with faster photocycles. Hence, photocurrent intensities for the WT and C303A were measured during continuous light illumination for 100 ms and were plotted against power density of the excitation light (Fig. 5f). As a result, the C303A mutant with the slower channel closing rates exhibited a lower EC_50_ value than that of the WT. Additionally, when compared to another BCCR, ChRmine, which has a slower closing rate than that of the HulaCCR1 WT (Fig. S4d), the trend that longer closing rates lead to lower EC_50_ values was maintained (Fig. 5f).

Previous studies on BCCRs suggested that amino acid residues on the intracellular side are also crucial for the gating mechanism. In *Gt*CCR2, a mutant in which D98 (equivalent to D96 in *Hs*BR) in TM3 was mutated to asparagine showed no significant channel current [40]. The effect of an identical mutation has been investigated in other BCCRs, including ChRmine [33] and *Hc*KCR1 [41], where the motif residues DTD are changed to DTN. This resulted in a decrease in cation channel activity. Moreover, it has been previously suggested for *Gt*CCR2 that proton transfer from D98 (corresponding to D96 in BR) to an unknown cytoplasmic residue and back govern the opening and closing of the channel, respectively [40]. We notice that BCCRs have characteristic well-conserved acidic amino acids on the C-terminal side of TM7 (Fig. S3). We thus hypothesized that a potential proton exchange between the third motif residue (D) in TM3 and acidic amino acid residues at the cytoplasmic side of TM7 in BCCRs plays a critical role in the channel gating. To test this hypothesis, we generated mutants corresponding to these residues in HulaCCR1: D187N in TM3, and E319Q, D320N, and E322Q, in TM7, and analyzed their photocurrents (Figs. S5a–c). The D187N and D320N mutants exhibited low expressions in the plasma membrane probably due to a significant decrease in protein stability, resulting in no observable photocurrents (Fig. S5c). Whereas E319Q exhibited photocurrent intensity and opening/closing rates similar to that of the WT (Figs. S5b and c), a decrease in the closing rate after 200-ms illumination was indeed observed for E322Q (17 ± 2 ms, 15 ± 1 ms, and 38 ± 2 ms for the WT, E319Q, and E322Q at −60 mV holding potential, respectively, *n* = 5, Fig. S5b).

Next, we compared photocurrent signals upon nanosecond laser-pulse illumination to precisely determine the difference in the time constants of opening and closing (Figs. S5d and e). Additionally, to clarify the relationship between the proton transfer reaction and channel dynamics, the kinetic isotope effect (KIE) in heavy water (D_2_O) was investigated by calculating a ratio between the time constants with D_2_O and H_2_O solutions with respect to the channel opening and closing kinetics. If a proton transfer is the rate-limiting elementary reaction step, the reaction rate decreases in D_2_O, thereby exhibiting a significant KIE value [42–44]. For the HulaCCR1 WT and E322Q, the time courses of photocurrents at membrane potentials of −60 mV and 40 mV are shown in Fig. S5d. The photocurrent kinetic traces of E322Q were fitted with a quadruple-exponential function, similar to the WT, to determine the time constants of channel opening and closing (Fig. S5e). By calculating the ratio of closing time constants in D_2_O to those in H_2_O, the KIE was determined to be 1.61 ± 0.06 for the WT and 1.18 ± 0.03 for E322Q, respectively. Hence, E322Q showed a smaller KIE for channel closing compared to that of the WT. Unfortunately, information about D187, which is homologous to *Hs*BR D96 and thought to be involved in the proton transfer with E322, could not be obtained due to the lack of significant photocurrents for D187N. However, considering the proton transfer mechanism proposed in the previous study for *Gt*CCR2, where protonation of the cytoplasmic D in TM3 controls channel closing, we hypothesized that a similar mechanism might occur in HulaCCR1 and the proton transfer from E322 to D187 governs the channel closing there as well.

### Long N/C-terminal extension of HulaCCR1

One of the prominent structural features of HulaCCR1 is the long N-terminal extension. A careful comparison of HulaCCR1 to other BCCRs, revealed that the first 12 residues of the expressed construct might not belong to the ORF. Expression and photocurrent of the N-terminally truncated variant (13–428) were recorded and those were similar to the full-length protein (1–428) (Figs. S6a and b). Even excluding the first 12 residues from consideration, there are still 79 residues before the first helix of the rhodopsin domain, which exceeds the length of this region in any other BCCR. The N-terminus harbors a distinctive amino acid sequence, which is predicted as signal peptides by signal peptide prediction methods. The exact cleavage site varied depending on the software used: SignalP 6.0 [45] predicted it to be between residues 43 and 44, while TOPCONS [46] predicted the cleavage site between 30 and 31. Similar signal peptides were found in other members of the mgCCR1–3 clade. To investigate the role of the N-terminal extension in HulaCCR1, two truncated variants were constructed: variant I without the predicted signal sequence region (residues 41–428) and variant II lacking most of the N-terminal region (residues 72–428).

No stop codon was observed in the 1285-nt contig encoding HulaCCR1, yet the 108 residues of the C-terminal extension were available for analysis. Similarly to other BCCRs, no homology could be detected to known functional domains in the C-terminal extension of HulaCCR1, but the structure modeled using AlphaFold2 exhibited two additional helices (Fig. 6a). Based on the structural model, two C-terminal truncations: variants III and IV, were produced by truncating immediately after the predicted TM7 helix (residues 1–341) and after the first additional helix after TM7 (residues 1–376), respectively. Patch-clamp measurements were conducted for these four truncated variants (I–IV).

**Fig. 6.**
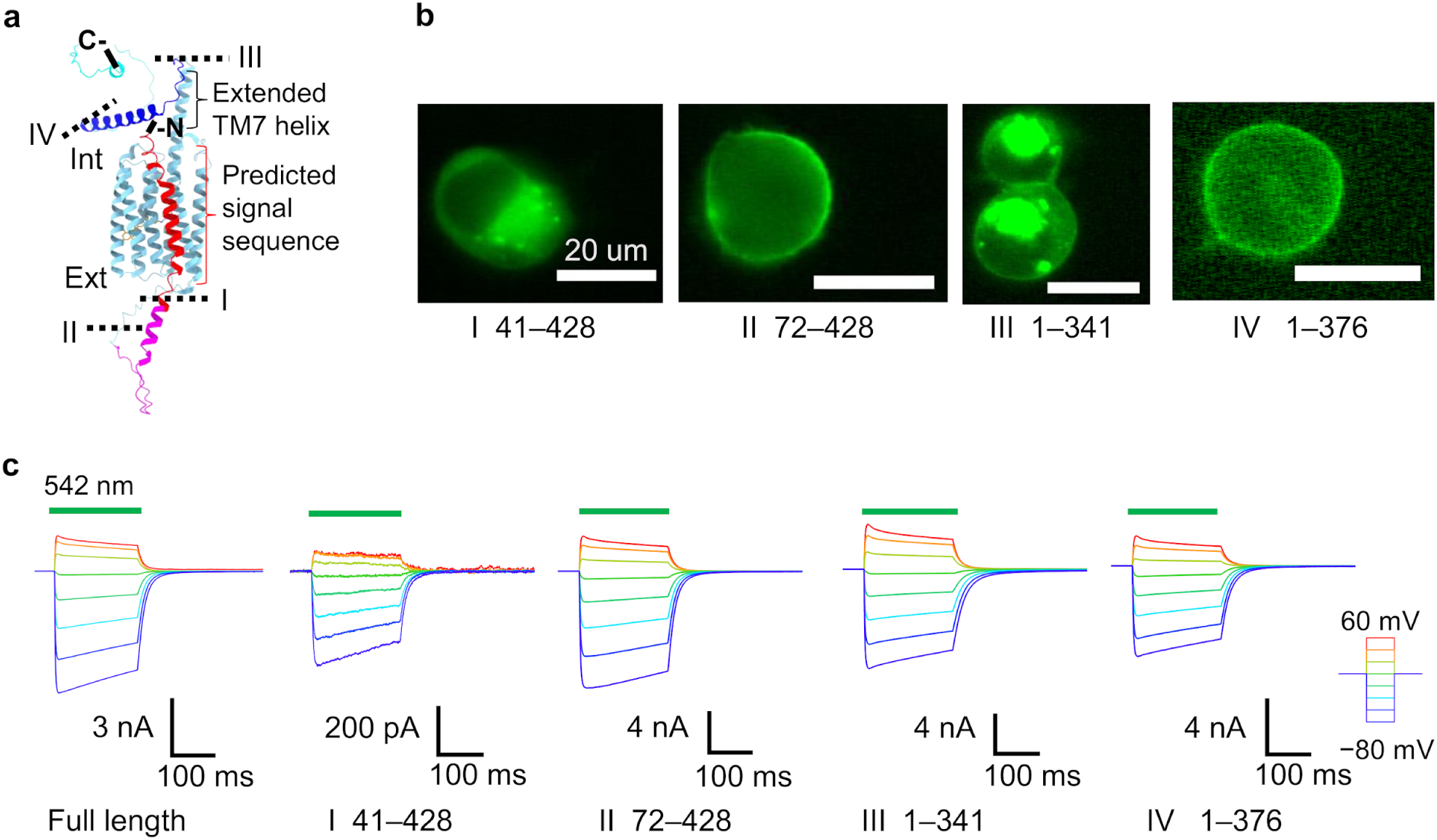
The long N/C-terminal extensions of HulaCCR1. **a** Truncated sites shown in the AlphaFold2 predicted structural model of HulaCCR1. **b** Fluorescence images of truncated variants expressed in ND7/23 cells. Scale bars: 20 μm. **c** Representative photocurrent traces of truncated variants recorded with standard pipette and extracellular solutions.

Based on the fluorescence images (Fig. 6b), truncated variant I (residues 41–428) exhibited significantly decreased plasma membrane trafficking in cells. When cells showing strong fluorescence were used, no significant photocurrent was obtained. However, significant photocurrents were observed with cells showing relatively lower fluorescence intensity, indicating that mild protein expression improves membrane trafficking of the protein (Fig. 6c). The shape of photocurrents of the truncated variant I did not differ from that of the full-length protein. Hence, the N-terminal extension appears to actually function as a signal peptide enhancing the membrane trafficking of HulaCCR1. Intriguingly, truncated variant II, which has a further truncated N-terminus (residues 72–428), membrane translocation was restored, showing an expression pattern and photocurrents similar to those of the full-length protein. We conclude that the region 41–71 between the predicted secretion signal and the rhodopsin domain appears to reduce the localization in the plasma membrane. However, the decreased membrane localization is compensated by the secretion signal peptide (residues 1–41) in the full-length protein. The region spanning the residues 41–71 is predicted by AlphaFold2 to have a disordered structure and is not conserved even among mgCCR1-like ChRs.

For the C-terminal truncated variants, truncated variant III, which is composed of the residues 1–341 and truncated immediately after TM7, showed an increase in fluorescence intensity inside the cells (Fig. 6b), indicating a decreased membrane localization compared to that of the full-length HulaCCR1. Truncated variant IV, in which a predicted disordered region beyond an additional cytoplasmic helix was removed, exhibited higher membrane localization than that of truncated variant III. The photocurrents of truncated variants III and IV were similar to that of the full-length HulaCCR1, indicating that the C-terminal extended region is not as essential as the N-terminal one (Fig. 6c).

### HulaCCR1 is able to drive neuronal firing

Our findings suggest that HulaCCR1 exhibits cation conductance and ion selectivity similar to those of *Gt*CCR4 and ChRmine, indicating its potential as an optogenetic tool to induce neuronal firing using light as other CCRs. To assess the practical utility of HulaCCR1 as an optogenetic tool, primary cultures of rat hippocampal neurons were transfected with HulaCCR1, and its expression was confirmed based on the fluorescence signal (Fig. 7a). HulaCCR1 was well expressed throughout the neuronal cells, including the axon. Patch-clamp recordings were performed on neurons expressing HulaCCR1 (Fig. 7b), and upon 200 ms light stimulation, neuronal firings were elicited. Additionally, even with 5 ms light stimulation at 10 Hz, continuous neuronal firing was observed, demonstrating HulaCCR1’s ability to control neuronal activity through light stimulation. This is likely facilitated by the relatively high conductance of HulaCCR1 to sodium.

**Fig. 7.**
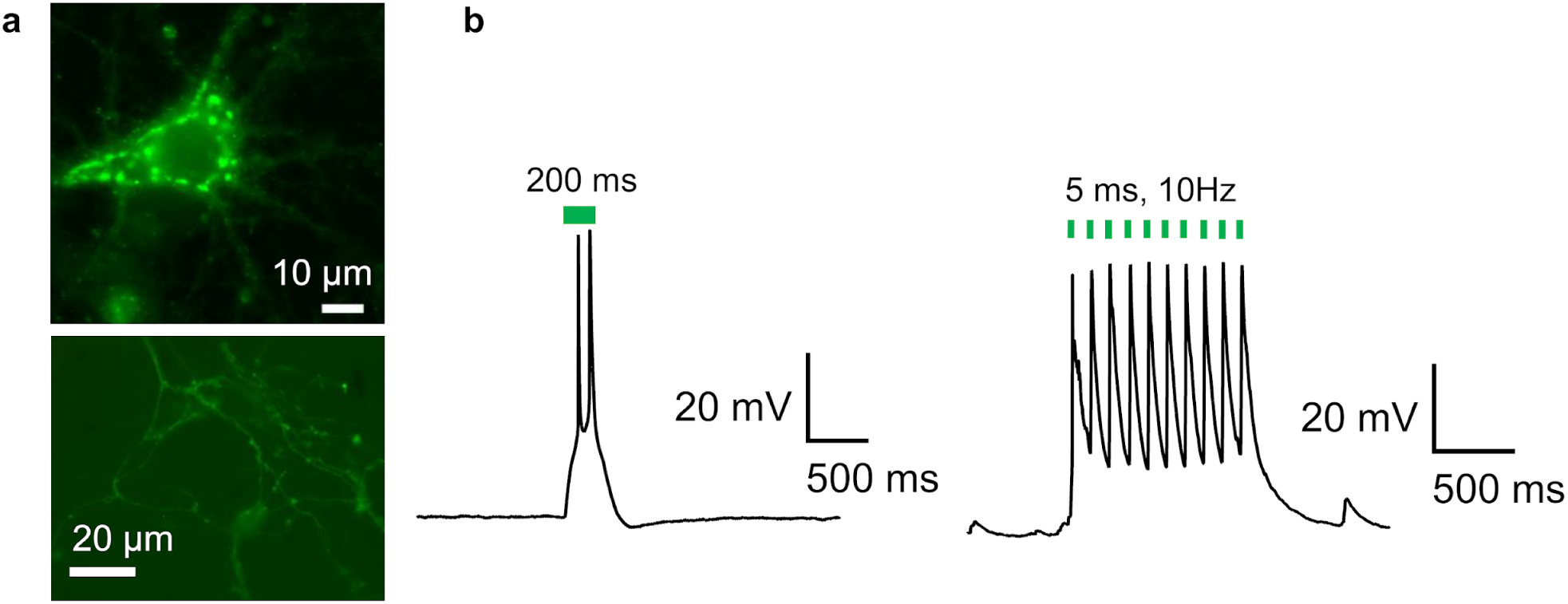
Light stimulation of hippocampal neurons expressing HulaCCR1. **a** Fluorescence images of the cell body (top) and the axon (bottom) of a hippocampal neuron expressing HulaCCR1. **b** Voltage traces showing depolarization and spikes of the neuron in response to 200 ms green (542 nm) light stimulation (left) and to 10 Hz stimulation with 5 ms pulses (right).

## Discussion

In this study, HulaCCR1 was identified as a new CCR featuring the ETD motif. Considering cation selectivity, *P*_Na+_/*P*_H+_ was previously estimated for *Cr*ChR2 and several other CCRs. While HulaCCR1 exhibits a *P*_Na+_/*P*_H+_ value of (1.5 ± 0.1) × 10^−5^, *P*_Na+_/*P*_H+_ of *Cr*ChR2 is approximately 1 × 10^−6^ [8]. *Gt*CCR4, with a value of about 5 × 10^−5^, has been reported as a CCR with low proton selectivity [32], suggesting that HulaCCR1 also has relatively low proton selectivity like *Gt*CCR4 among CCRs. The *P*_X+_/*P*_H+_ values obtained for HulaCCR1 do not differ significantly between the different alkali-metal monovalent cations, indicating similar permeability levels for all of them. The photocurrents derived from Cs^+^ in HulaCCR1 do not significantly differ from that of Na^+^, whereas the photocurrents from Cs^+^ are approximately half of that from Na^+^ in *Gt*CCR4 [32]. Hence, HulaCCR1 likely possesses a relatively large channel pore facilitating the transport of large monovalent cations. This is further supported by the fact Li^+^, which is strongly hydrated in solution [47], showed permeability comparable to other monovalent cations. Considering the expected large pore size capable of efficient Cs^+^ transport, it is likely that partially hydrated Li^+^ is stabilized within the pore rather than undergoing complete dehydration. mgCCR1 despite sharing the ETD motif with HulaCCR1 and having a 48% amino acid sequence identity, exhibits relatively high proton selectivity compared to HulaCCR1 [22]. The significant difference in ion selectivity despite the relatively high amino acid sequence identity is intriguing. A comparison of predicted structures using AlphaFold2 and cavities potentially involved in channel gating did not suggest significant differences in the size of channel pore (Fig. S6). Hence, although it is anticipated that the structural changes in the open state of HulaCCR1 are larger than that in mgCCR1, leading an opening of a larger channel that is capable of conducting large monovalent cations, further investigation is needed to clarify the reason of different cation selectivity between HulaCCR1 and mgCCR1.

Considering the role of the first motif residue E, it was found to influence the absorption spectrum and gating (Figs. 5b–d). In the E176D mutant, the time constants of channel opening and closing were altered compared to the WT, and regarding the fast channel closing component, the closing time constant was 1.8 and 1.3 times slower than that of the wild type at a +40 mV and a −60 mV membrane potential, respectively, indicating the difference in the closing rate between HulaCCR1 E176D and the WT becomes larger at positive membrane potential. Hence, in the WT, E187 appears to enhance inward rectification. Indeed, when observing the photocurrent upon 200 ms illumination, the positive photocurrent intensity in the WT is slightly suppressed compared to E176D (Fig. S4a). This suggests that the glutamate is one of the important residues defining the rectification properties of HulaCCR1.

Looking at the photocurrents of the C303A mutant involving the residue immediately before the retinal-binding lysine residue in TM7 at 200 ms light illumination, we find that the decay rate after light off is significantly slower compared to that of the WT (WT = 17 ± 2 ms, C303A = 130 ± 20 ms, at −60 mV, *n* = 5) (Fig. S4a). *Gt*ACR1, a member of the cryptophyte ACR family, also has a cysteine at this position, C237, and an analogous mutation, C237A, was reported to eliminate one of the two closing components and to show approximately 1.8 times slower closing rate for the remaining component [48]. However, the retardation effect of the corresponding mutation on the closing rate is much higher in HulaCCR1 (7.6 times slower), indicating that C303 is likely to influence the closing rate through a mechanism different from that of *Gt*ACR1. Since C303 is located close to the retinal Schiff base, the mutation of this residue may affect the photochemical reactions around the chromophore. Nevertheless, we observed no significant red shift in HulaCCR1 C303A based on the action spectra of the photocurrent (Fig. S4c), contrasting with the red shift in C237A of *Gt*ACR1 [48]. Hence, the interaction between the retinal chromophore and C303 in HulaCCR1 is different from that in *Gt*ACR1, and the effect on the closing process differs even with the identical mutation.

The N-terminal extension, outside the seven TM helices constituting the rhodopsin domain, contains a sequence predicted as a secretion signal sequence, and our results suggest that this secretion signal sequence rescues the decrease in membrane translocation caused by the sequence between residues 41 and 71. mgCCR1 has a similarly long N-terminal extension, although it is shorter and demonstrates low homology to that of HulaCCR1. Analogously, the N-terminal extension of mgCCR1 is predicted to have signal peptide cleaved between positions 24 and 25 (according to SignalP 6.0). Hence, it is anticipated that a similar mechanism maintains protein membrane translocation in mgCCR1 as well. Regarding the extension at the C-terminus, although not as prominent as the N-terminus, a slight decrease in membrane translocation compared to the full-length protein was observed in variant III (residues 1–341) (Fig. 6b). Hence, the C-terminal region may contribute to maintaining the stability of proteins in the membrane by interacting with the cytoplasmic surface of the rhodopsin part. The long C-terminal extensions of green algal CCRs, prasinophyte ACRs and labyrinthulean ACRs have been found to contain response regulator-like domains [49] as well as additional conserved domains of unknown function [49, 50], but we could not identify any such functional domains in HulaCCR1 or other BCCRs. It has been previously reported that multiple arginines within the C-terminal extension interact with the rhodopsin part and regulate the closing rate in the channelrhodopsin *Kn*ChR derived from a green alga *Klebsormidium nitens* [51]. HulaCCR1 also has several arginines between residues 352 and 358 that were truncated in variant III. Moreover, a slightly slower decay rate after light off was observed in variant III compared to that of the WT upon 200 ms light illumination (WT = 17 ± 2 ms, variant III = 21 ± 1 ms, at −60 mV, *n* = 5), exhibiting behavior similar to *Kn*ChR. Hence, it is conceivable that in structural stabilization in the membrane is facilitated by electrostatic interactions between the C-terminal extension and the cytoplasmic side of the rhodopsin domain mediated by the arginines, potentially influencing gating kinetics.

## Materials and Methods

### Sampling, RNA extraction, and metatranscriptome assembly

Peat lake sampling was performed on February 23^rd^, 2021 in The Hula Nature Reserve, Israel (33°04’33.8”N 35°36’40.8”E). 20 L of water from the surface and 0.5 m depth was filtered through a 50 µm mesh net to remove large particles, filtered onto a GF/D filter (Whatman), and flash-frozen using liquid nitrogen on-site.

Total RNA was extracted from frozen filters, corresponding to ∼5 L per sample (RNeasy PowerSoil Total RNA Kit; Qiagen). The RNA was tested with Bioanalyzer, and PolyA-selected libraries were constructed and sequenced at the Weizmann Institute of Science, on the NovaSeq platform (300 cycles, PE).

*De-Novo* assembly was done post trimming (Trimgalore; v. 0.6.6)[52], Cutadapt v. 3.4 [53] with MEGAHIT (v. 1.2.9) [54] and then clustered to reduce redundancy with cd-hit-est v. 4.8.1 [55]. An initial rhodopsin screening was done with HMMsearch (v. 3.3.2) [56] with the microbial rhodopsin profiles from https://github.com/BejaLab/RhodopsinProfiles.

### Bioinformatic analyses

mgCCR1, mgCCR2, mgCCR3, HulaCCR1, and other members of the mgCCR1-3 clade from the Hula lake were used to search for additional datasets from JGI Integrated Microbial Genomes and Microbiomes (IMG) [57] containing members of the clade. An initial search for such datasets was performed using two strategies: a large collection of proteins assigned in IMG to PFAM family pf01036 from a broad set of environmental assemblies was searched using blastp from NCBI blast package v. 2.12.0+ [58] and a narrower collection of entire aquatic eukaryotic metatranscriptome assemblies was searched directly using tblastn. For the matching assembled and unrestricted datasets, as well as the lake Hula assembly, the entire assemblies were searched against a database of rhodopsin domains of BCCRs and curated representatives of the mgCCR1-3 clade using blastx with an e-value threshold of 1e-5. The contigs were assigned to the mgCCR1-3 clade if they had best hits to one of the mgCCR1-3 clade representatives with at least 25 amino acid alignment length and a minimal identity of 65%. ORFs in the matching transcript fragments and exons in the genomic fragments were predicted manually based on multiple sequence alignment for each subclade. To resolve phylogenetic relationships between the ChRs of the mgCCR1-3 clade, the most complete representative sequences were chosen and combined with other members of the stramenopile CCR family. The protein sequences were aligned with mafft v. 7.526 [59] and trimmed to include the region between the transmembrane helices 1–7 (A–G) using the structure of *Hc*KCR1 (PDB: 8H86) [41] as a template. Maximum likelihood phylogeny was reconstructed with RAxML v. 8.2.13 [60] under the PROTCATLG model and with 1000 fast bootstrap replicates. The rest of the environmental sequences were placed on the resulting tree using pplacer v. 1.1.alpha19 [61] by adding them to the alignment with mafft. The phylogenetic placements were resolved with gappa v. 0.8.0 [62].

Analysis of retrotransposons associated with the ChR-possessing genomes was performed as follows. Regions upstream and downstream of the start and stop codons of the genes from the four freshwater ChR subclades were extracted and sufficiently long sequences (≥150 nt) were clustered with CD-HIT v. 4.8.1 [63] at 90% identity level and used as a database to search for related matches in the same environmental assemblies with blastn. Contigs matching the regions flanking the rhodopsin genes with E value ≤ 1e-12 were extracted and searched for (retro)transposons using TransposonPSI v. 1.0.0 (https://transposonpsi.sourceforge.net/) with an E value threshold of 1e-10. Related retrotransposons were searched for using the resulting protein sequences of the *pol* genes using blastp in three eukaryotic databases: EukProt v. 3 [64], Tara Oceans Single-Cell and Metagenome Assembled Genomes (SMAGs) [65] and single-cell assembled genomes from bioproject PRJNA379597 [24]. Matches passing the empirical bitscore threshold of 250 were clustered with cd-hit at 62% identity level, combined with non-redundant query *pol* genes (clustered at 95% protein identity level), aligned with mafft (automatic mode selection) and trimmed with trimAl v. 1.4.1 (-automated1 method) [66]. Trimmed sequences longer than 200 amino acids were taken to phylogenetic reconstruction with RAxML under the PROTCATLG model, with 1000 fast bootstrap replicates. The shorter sequences were placed on the resulting tree with pplacer.

For the analysis of the eukaryotic community composition, SSU rRNA fragments were extracted from the assemblies using metaxa v. 2.2.3. Taxonomy was assigned to the resulting sequences using sintax from usearch v. 11.0.667 [67] with Protist Ribosomal Reference (PR²) database v. 5.0.0 [68], with the default cut-off 0.8. Only matches to Eukaryota excluding animals were considered. For SSU rRNA phylogenetic tree of the kathablepharids, non-chimeric kathablepharid sequences and sequences from chosen outgroups longer than 1400 nt were extracted from PR², clustered at 99% identity level with cd-hit and combined with SSU sequences from the deep-sea kathablepharid SAGs and one of the complete kathablepharid SSU sequences from the assemblies containing mgCCR1-3 genes. The phylogenetic analysis was performed by aligning the sequences with mafft, trimming the alignment with trimAl (-automated1 method) and reconstructing the phylogeny using RAxML under the GTRCAT model, with 1000 fast bootstrap replicates.

The snakemake workflow implemented for the bioinformatic analyses is available at https://github.com/BejaLab/hulaccr.

### DNA constructs

Coding sequence (Table S1) of full-length HulaCCR1 and ChRmine were cloned into pCMV3.0-enhanced fluorescent yellow protein (EYFP) vector between EcoRI and BamHI sites with a Kir2.1 membrane trafficking signal, EYFP, and an ER-export signal [69] fused at the C-terminus of HulaCCR1. For point mutations, the QuikChange site-directed mutagenesis method (Agilent Technologies, CA) was employed according to a manufacturer standard protocol. The truncated mutants were PCR-based cloned using EcoRI-BamHI restriction sites into the pCMV3.0-EYFP vector. Primers used in DNA work are listed in Table S2.

### UV–visible spectroscopy

Cell culture and transfection for UV–visible spectroscopy were conducted as described elsewhere [70]. Briefly, COS-1 cells (cell number: JCRB9082; Japanese Collection of Research Bioresources Cell Bank, Osaka, Japan) were transfected by the polyethyleneimine method [71]. Cells were supplemented with 2.5 µM all-*trans*-retinal (ATR) on the following day and harvested after 48 h of transfection. The cell membranes were solubilized with a phosphate buffer (pH 7.0) containing (in mM) 66.5 phosphate, 133 NaCl with 3% *n*-dodecyl-β-D-maltoside. After centrifugation (21,600 × g, 20°C, 5 min), the supernatant was subjected for the measurement of UV–visible absorption spectra using a spectrophotometer (V-730, JASCO, Japan). Rhodopsin was bleached by hydroxylamine (50 mM) and illumination with visible light from a 1-kW Xe lamp (MAX-303, Asahi Spectra, Japan) through a long-pass filter (Y52, AGC Techno Glass, Japan). Difference spectra were calculated by subtracting post-illumination spectra from the pre-illumination spectrum. To correct for spectral changes derived from non-rhodopsin components in the sample, the corresponding spectra of untransfected cell membranes that underwent the same treatment as the sample were subtracted from the sample spectra.

### Cell culture and transfection for electrophysiology

ND7/23 cells were grown in Dulbecco’s modified Eagle’s medium (D-MEM, FUJIFILM Wako Pure Chemical Co., Japan) supplemented with 5% fetal bovine serum (FBS) under a 5% CO_2_ atmosphere at 37°C. ND7/23 cells were attached onto a collagen-coated 12-mm coverslips (IWAKI, cat. 4912-010, Japan) placing in a 4-well cell culture plate (SPL Life Sciences, cat. 30004, Korea). The expression plasmids were transiently transfected in ND7/23 cells using Lipofectamine 3000 transfection reagent (Thermo Fisher Scientific Inc., MA). Seven to eight hours after the transfection, the medium was replaced with D-MEM containing 10% horse serum (New Zealand origin, Thermo Fisher Scientific Inc., MA), 50 ng mL^−1^ nerve growth factor-7S (Sigma-Aldrich, MO), 1 mM N^6^,2’-O-dibutyryladenosine-3’,5’-cyclic monophosphate sodium salt (Nacalai tesque, Japan), 1 μM cytosine-1-β-D(+)-arabinofuranoside (FUJIFILM Wako Pure Chemical Co., Japan), and 2.5 μM all-*trans*-retinal. Electrophysiological recordings were conducted at 1–3 days after the transfection. The transfected cells were identified by observing the EYFP fluorescence under an up-right microscope (BX50WI, Olympus, Japan).

Hippocampi were isolated from postnatal day 1 (P1) Wistar-ST rats (by Sankyo Labo Service Co., Japan) and treated with papain dissociation system (Worthington Biochemical Co., NJ) according to the company protocol. Briefly, dissected hippocampi were treated with papain for 1 h at 37°C. The dissociated cells were washed with Earle’s Balanced Salt Solution (EBSS) supplemented with ovomucoid protease inhibitor with bovine serum albumin. The dissociated cells were resuspended in culture medium (Neurobasal A containing 2% B-27 plus (Thermo Fisher Scientific Inc., MA)). Approximately 150,000 cells were plated on 12-mm poly-L-lysine-coated glass coverslips (IWAKI, cat. 4912-040, Japan) in the 4-well cell culture plate. Transfection was done by adeno-associated virus (AAV), and the plasmid and virus packaging were synthesized by Vectorbuilder Inc. The plasmid pAAV vector has a CaMKIIa promotor containing the sequence of HulaCCR1, C-terminally fused to the Kir2.1 membrane trafficking signal followed by EYFP and the ER export signal. For transfection, AAV8 was added to each well (4–8 × 10^8^ genome copies well^−1^) 7 days after plating. The electrophysiological recordings were performed 15–22 days after transfection.

### Electrophysiology

All experiments were carried out at room temperature (20–22°C). Currents were recorded using an EPC-8 amplifier (HEKA Electronic, Germany) under a whole-cell patch clamp configuration. The data were filtered at 1 kHz, sampled at 50 kHz (Digidata1440 A/D, Molecular Devices Co., CA) and stored in a computer (pClamp11.1, Molecular Devices). The standard internal pipette solutions for the whole-cell voltage clamp recordings from the ND7/23 cells contained (in mM) 119 KF, 10 KOH, 5 Na_2_EGTA, 14 HEPES, 0.0025 ATR (pH 7.4 adjusted with HCl). The standard extracellular solution contained (in mM): 138 NaCl, 3 KCl, 2.5 CaCl_2_, 1 MgCl_2_, 10 HEPES, 4 NaOH, and 11 glucose (pH 7.4 adjusted with HCl). In the ion selectivity measurement, extracellular solution contained (in mM) 146 XCl (X = Na^+^, K^+^, Li^+^, Rb^+^, Cs^+^), 6 N methyl-D-glucamine (NMG), 2.5 CaCl_2_, 10 HEPES, 11 glucose (pH 7.4 adjusted with HCl). For the divalent cation selectivity, extracellular solution contained (in mM) 70 XCl_2_ (X = Ca^2+^, Mg^2+^), 6 NMG, 2.5 CaCl_2_, 10 HEPES, 11 glucose (pH 7.4 adjusted with HCl). The NMG-extracellular solution contained (mM) 146 NMG, 146 HCl, 2.5 CaCl_2_, 10 HEPES, and 11 glucose (pH 7.4 adjusted with NMG). The NMG-pipette solution contained (in mM) 130 NMG, 90 glutamate, 6 (NMG)_2_EGTA, 50 HEPES, 2.5 MgSO_4_, 2.5 MgATP, 0.0025 ATR (pH 7.4 adjusted with H_2_SO_4_). The liquid junction potentials (LJPs) were calculated by pClamp 11.1 software (Table. S3) and estimated reversal potentials were compensated by calculated LJPs.

For whole-cell voltage clamp, illumination at 377 ± 25, 438 ± 12, 472 ± 15, 510 ± 5, 542 ± 13, 575 ± 12 or 643 ± 10 nm was carried out using a SpectraX light engine (Lumencor Inc., OR) controlled by pClamp 11.1 software. HulaCCR1 was illuminated through an objective lens (LUMPlan FL 40x, NA 0.80W, Olympus, Japan). For the standard photocurrent detection, the power of the 542-nm light was directly measured under a microscope using a visible light-sensing thermopile (MIR178 101Q, SSC Co., Ltd., Japan) and was adjusted to 13.3 mW mm^−2^. The action spectrum was measured at a holding potential of −40 mV at wavelengths with equivalent power density of 0.8 mW mm^−2^. Each action spectrum was estimated by the maximal amplitude of the photocurrent scaled by the light power density under the assumption of a linear relationship between the photocurrent intensity and the light power density.

For Laser-flash patch clamp experiment, a laser flash (3–5 ns) at 532 nm (Nd:YAG laser, Minilite II, Continuum, CA) was used [44]. For measuring the KIE on the gating dynamics, the internal pipette solutions were made of H_2_O or D_2_O solutions containing (in mM) 60 Na_2_SO_4_, 25 NMG, 60 mannitol, 5 EGTA, 10 HEPES, 2.5 MgSO_4_, 2.5 MgATP, and 0.0025 ATR adjusting pH to 7.4 with HCl or DCl, respectively. The extracellular solutions were made of H_2_O or D_2_O solutions containing (in mM) 133 NaCl, 3 KCl, 5 NMG, 10 HEPES, 5 CaCO_3_ (solved by HCl or DCl solutions), 2.5 MgSO_4_ and 11 glucose, adjusting pH to 7.4 with HCl or DCl, respectively. The pipette resistance was adjusted to 3–5 MΩ with a series resistance of 7–19 MΩ (WT; 12 ± 1, *n* = 9 for H_2_O; 16 ± 1, *n* = 9 for D_2_O, E176D; 9 ± 1, *n* = 9 for H_2_O, E322Q; 10 ± 1, *n* = 9 for H_2_O; 15 ± 1, *n* = 9 for D_2_O) and a cell capacitance of 30–90 pF (WT; 63 ± 4, *n* = 9 for H_2_O; 58 ± 3, *n* = 9 for D_2_O, E176D; 50 ± 2, *n* = 9 for H_2_O, E322Q; 43 ± 3, *n* = 9 for H_2_O; 44 ± 2, *n* = 9 for D_2_O). In every experiment, the series resistance was compensated for by 70%. Obtained signals had electrical noise from the laser system. Therefore, the noise signal recorded without light excitation was subtracted from raw photocurrent traces.

For whole-cell recordings in cultured hippocampal neurons, patch pipettes were filled with (in mM) 90 potassium gluconate, 30 KOH, 49 HEPES, 2.5 MgCl_2_, 2.5 MgATP, 5 Na_2_EGTA, and 0.0025 ATR, titrated to pH 7.4. The extracellular solution contained (in mM) 138 NaCl, 3 KCl, 2.5 CaCl_2_, 1 MgCl_2_, 11 glucose, and 10 HEPES, 2 kynurenic acid, titrated to pH 7.4.

## Acknowledgements

We thank Ariel Chazan, Shirley Larom, David Iluz, Said Abu-Ghosh, and the park rangers David Zaguri and Ido Shaked for help in sampling, and the Israel Nature and Parks Authority for the permit to sample in The Hula Nature Reserve. This work was supported by JSPS KAKENHI Grants-in-Aid (grant JP23K14151 to S.T., JP21H01875, JP23K18090, and JP23H04404 to K.I.), JST CERST (grant JPMJCR22N2 to K.I.), and MEXT Promotion of Development of a Joint Usage/ Research System Project: Coalition of Universities for Research Excellence Program (CURE) (grant JPMXP1323015482 to K.I.), the European Commission, under Horizon Europe’s research and innovation programme (Bluetools project, Grant Agreement No 101081957 to O.B.), the Israel Science Foundation (Research Center grant 3131/20 to O.B.), and the Nancy and Stephen Grand Technion Energy Program (GTEP). O.B. holds the Louis and Lyra Richmond Chair in Life Sciences.

## Supporting information

**Fig. S1.**
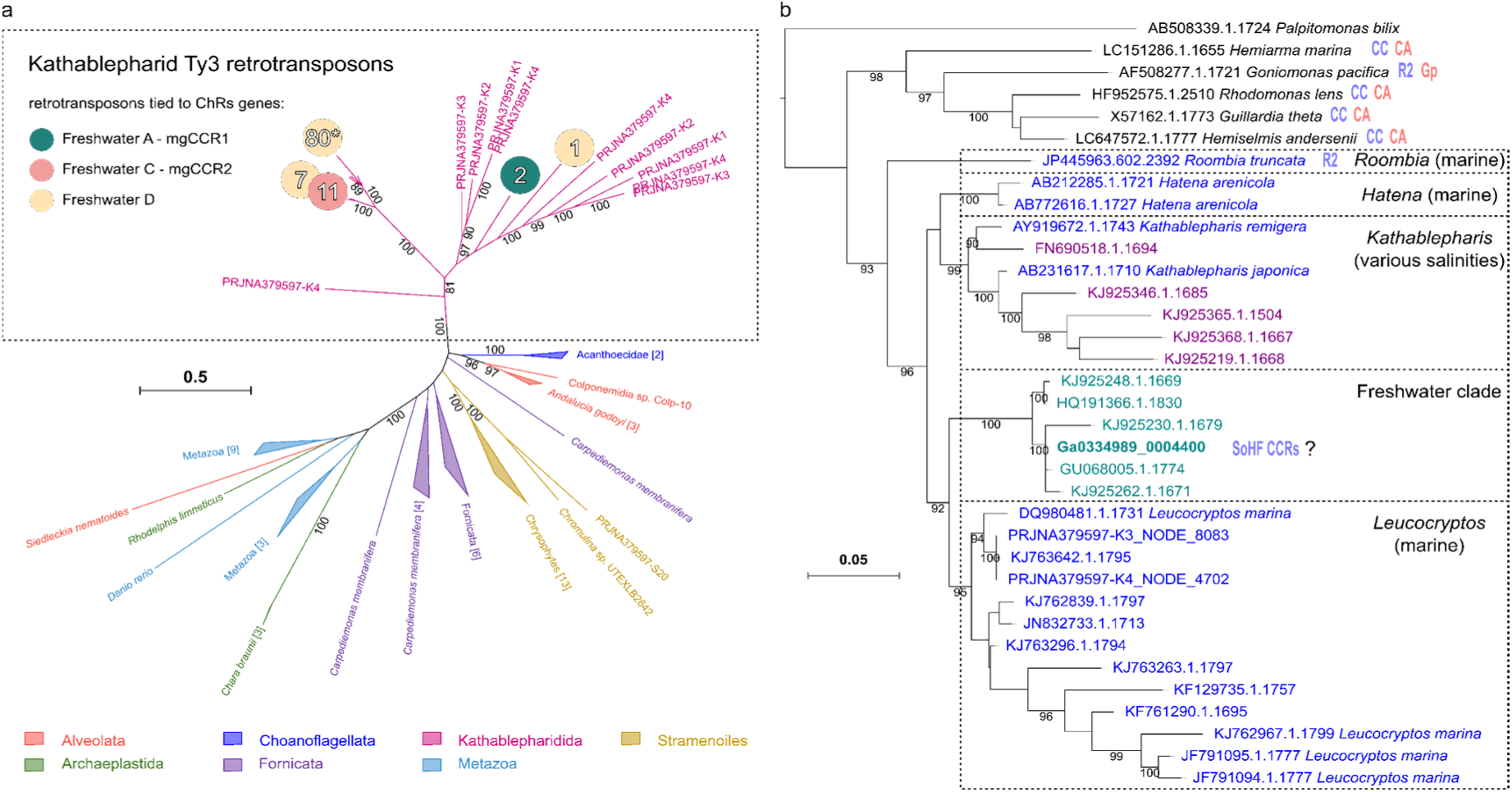
Evolutionary affinities of the uncultured eukaryotes possessing ChRs from the mgCCR1-3 clade. **a** Phylogenetic analysis of the Ty3/gypsy retrotransposons linked to genes coding for three freshwater subclades of the ChRs. Branches are labeled by the corresponding taxonomic group with the kathablepharid genomes represented by four SAGs K1-K4 (bioproject PRJNA379597). Branches containing *pol* genes from the metagenomic contigs are indicated with circles colored by the subclade of the ChR genes linked to them (see Materials and Methods for details). Numbers inside indicate the number of the individual contigs containing fragments of the corresponding retrotransposons. Asterisk indicates position of the contig Ga0335041_0096840 which contains a fragment of a retrotransposon upstream of a ChR gene. Numbers next to branches indicate fast bootstrap support values ≥80. **b** Phylogenetic tree of SSU rRNA genes from kathablepharids and other cryptists. Contig Ga0334989_0004400 is a complete gene representative of a dominant kathablepharid SSU type found in datasets with ChRs from the mgCCR1-3 clade. Labels are colored by the sample environment: blue — marine, cyan — freshwater, purple — estuarine. Red and blue abbreviated labels indicate the presence of ACR and CCR families in the corresponding species: CA — cryptophyte ACRs, CC — cryptophyte CCRs, Gp — GpACR1, R2 — mgR2-like ChRs, SoHF CCR — stramenopile (and other HF) CCRs. Numbers next to branches indicate fast bootstrap support values ≥90.

**Fig. S2.**
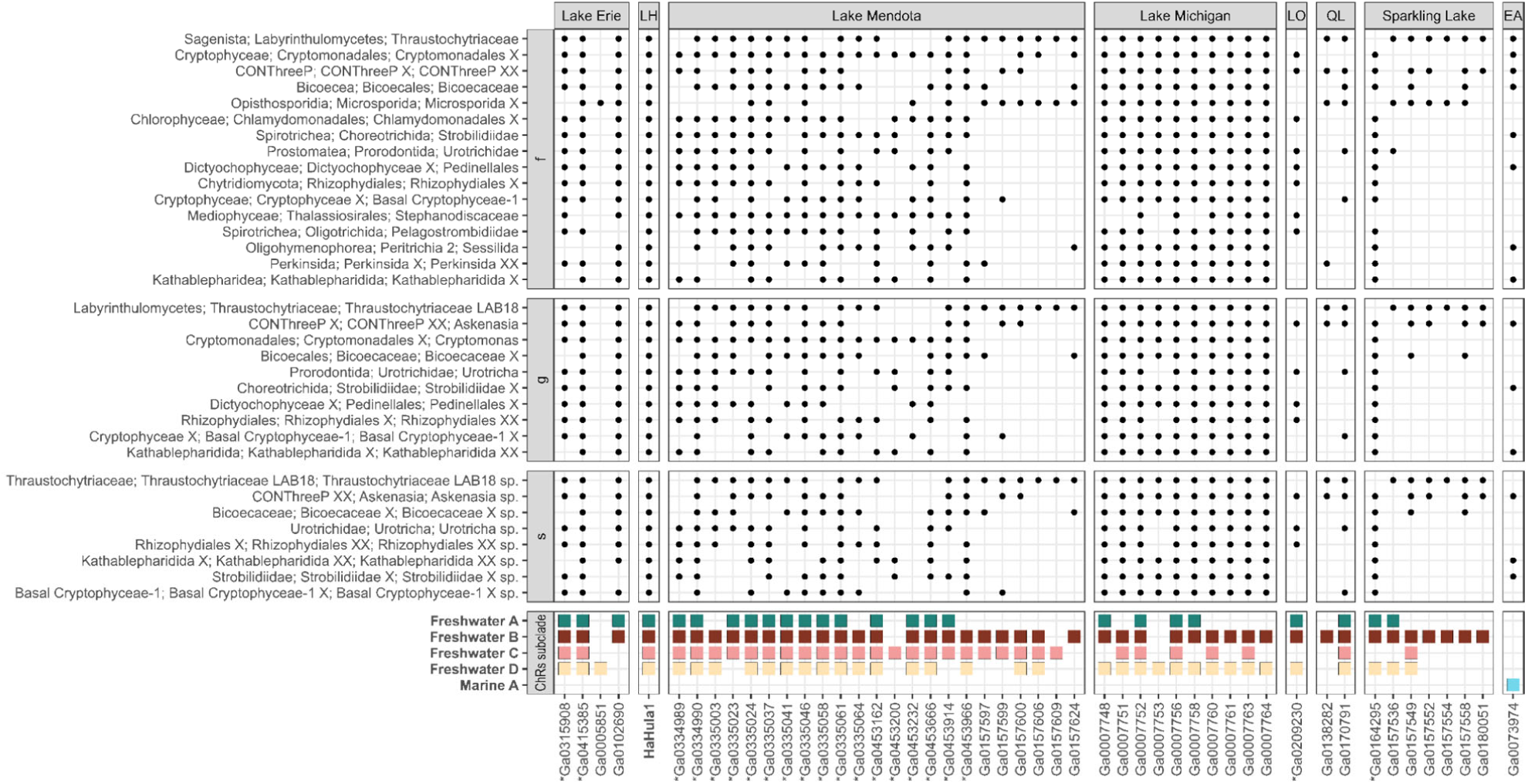
rRNA SSU taxonomic profiling of the environmental samples in which ChRs from the mgCCR1-3 clade were found. Taxa are grouped by rank as defined in the Protist Ribosomal Reference database at three levels: s, g and f. Shown are unicellular eukaryotic taxa present in at least 25 individual datasets at taxonomic levels. Datasets with too few rRNA contigs are not shown. Location abbreviations are as follows: LH — Lake Hula, LO — Lake Ontario, QL — Quebec Lakes, EA — equatorial Atlantic Ocean.

**Figure S3.**
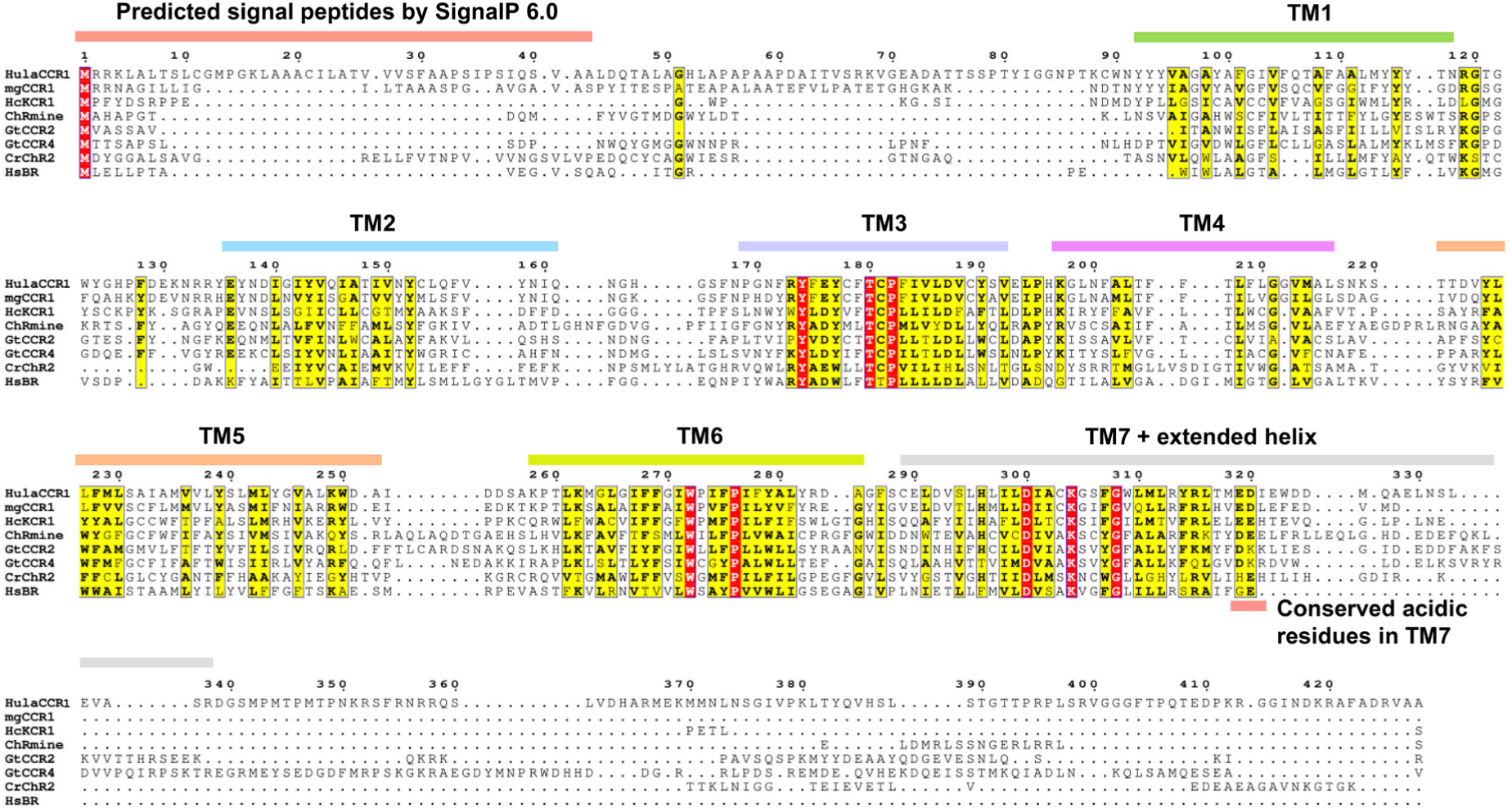
Sequence alignment of HulaCCR1 with representative channelrhodopsins and *Hs*BR. The sequence alignment was created using ClustalW [1] and ESPript 3 [2] servers. A predicted signal sequence, transmembrane helices, and conserved acidic residues in TM7 based on the AlphaFold2 structural model of HulaCCR1 are highlighted by colored bars.

**Figure S4.**
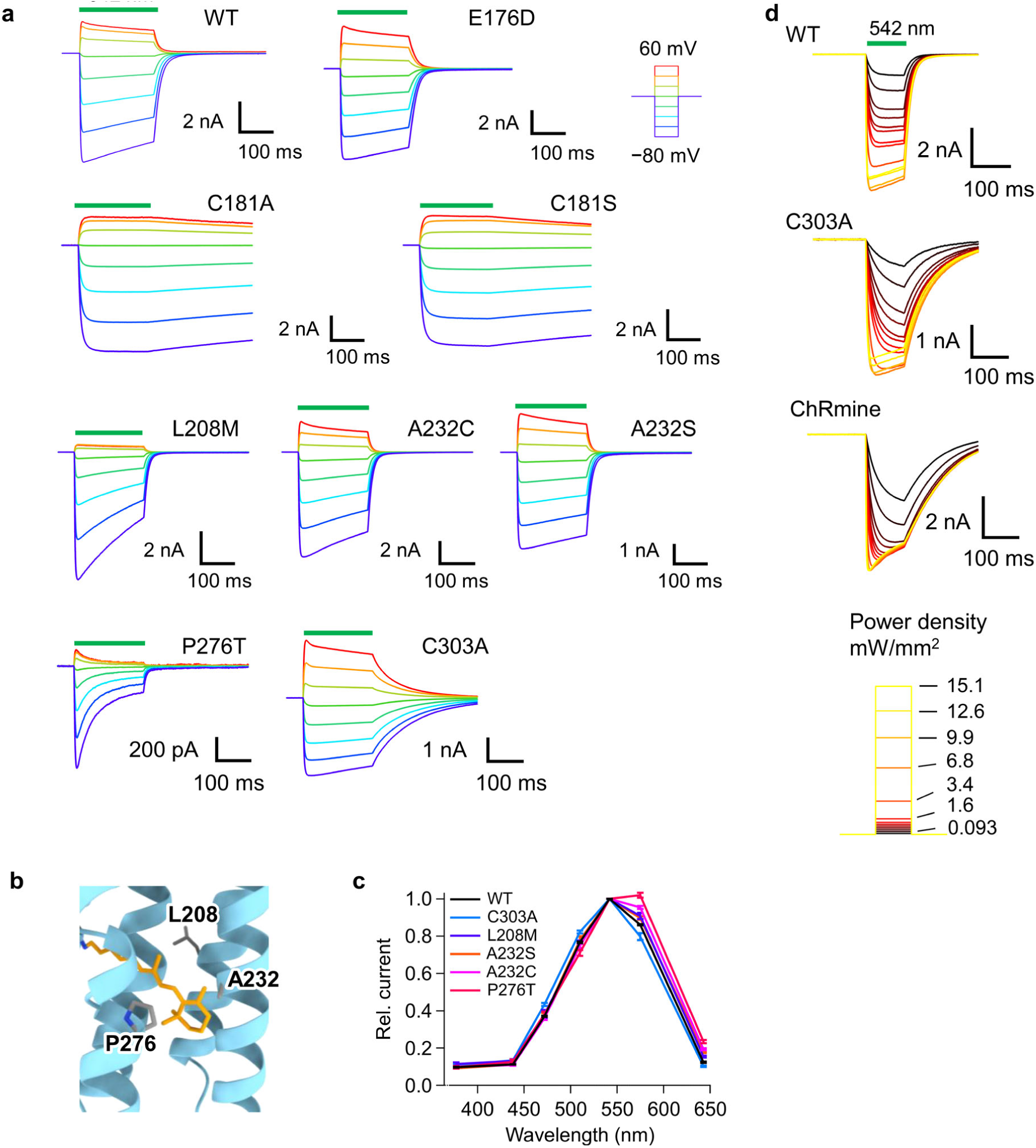
Photocurrents of HulaCCR1 mutants. **a** Representative photocurrent traces of the HulaCCR1 WT and mutants recorded with standard pipette and extracellular solutions. **b** Mutated residues close to the β-ionone ring side of the retinal. The retinal structure (orange) and the side chain of conjugated lysine are of *Hc*KCR1 (PDB ID: 8H86). **c** Normalized action spectra of the HulaCCR1 WT, C303A, L208M, A232S, A232C, and P276T (mean ± S.E., *n* = 5). **d** Light-power dependence of photocurrent traces of the HulaCCR1 WT, C303A, and ChRmine at −40 mV with standard pipette and extracellular solutions.

**Figure S5.**
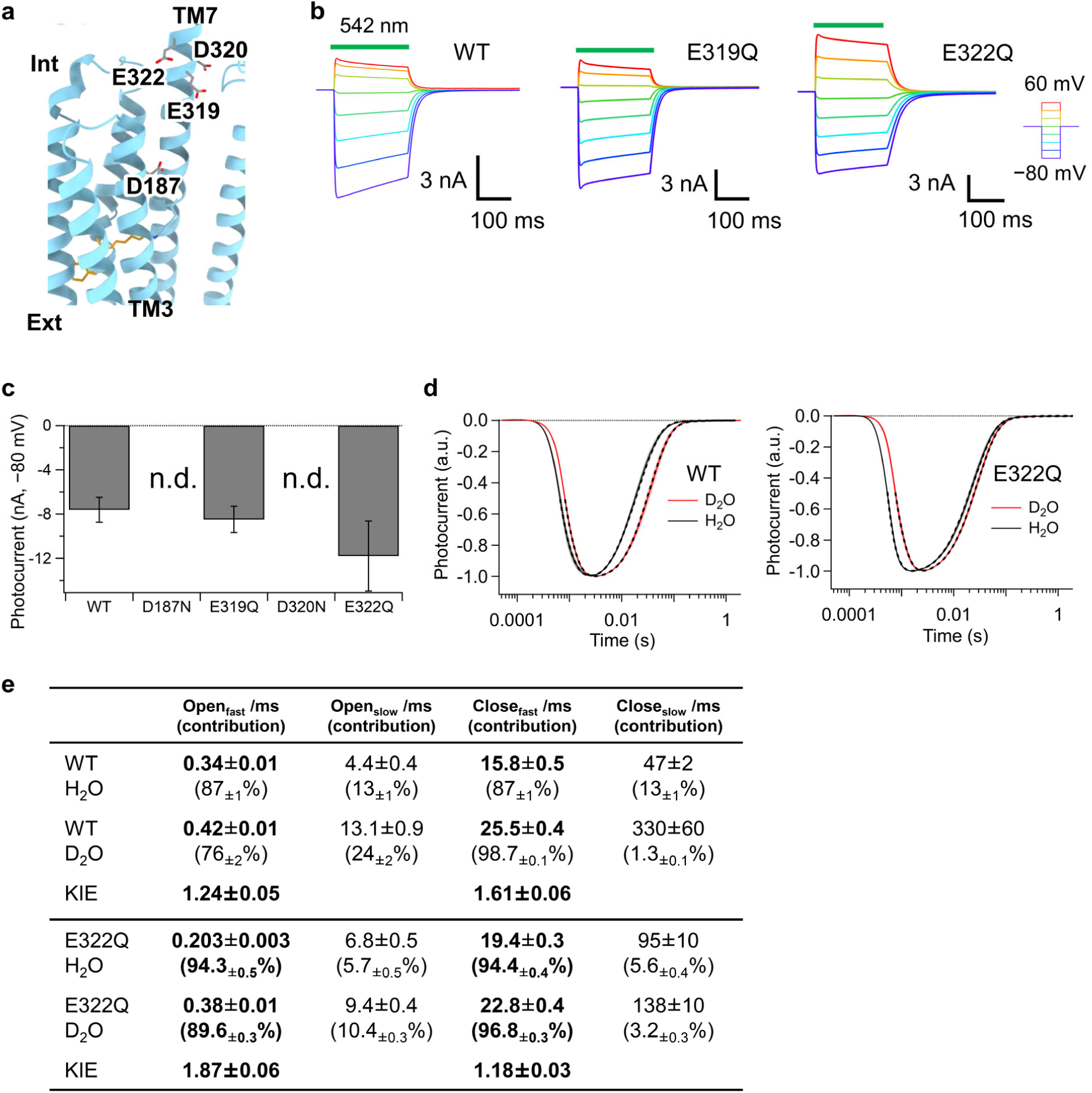
Effect of point mutations on intracellular acidic residues. **a** Mutated acidic residues located in the intracellular region. **b** Representative photocurrent traces of the HulaCCR1 WT, E319Q, and E322Q. **c** Mean photocurrent amplitude recorded at −80 mV (*n* = 5). **d** Photocurrent traces of the HulaCCR1 WT and E322Q upon nanosecond laser-flash excitation recorded in H_2_O/D_2_O solutions. Fitting curves are shown as broken lines. **e** Time constants of channel opening and closing of the HulaCCR1 WT and E322Q (mean ± S.E., *n* = 9)

**Figure S6.**
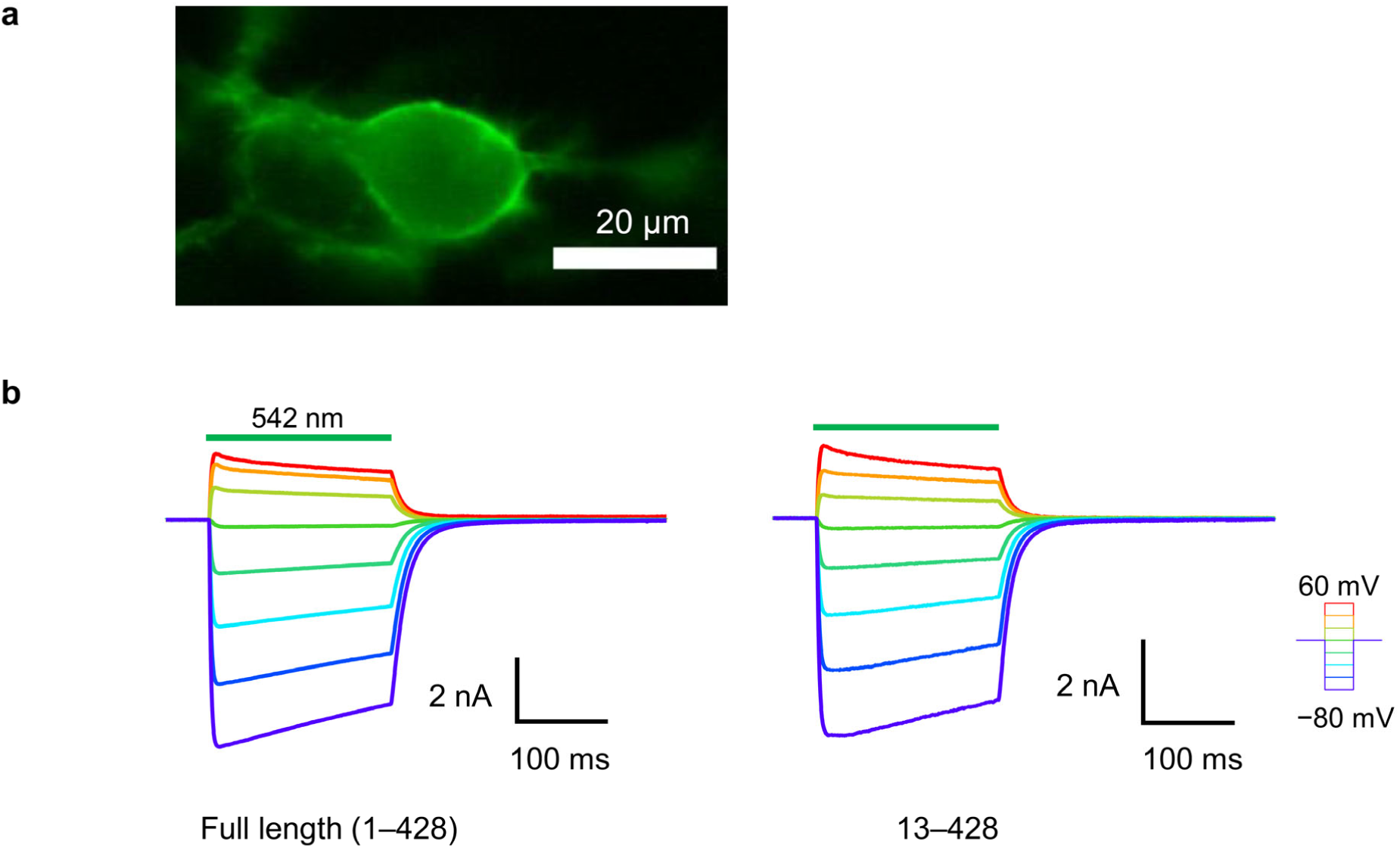
Comparison of the photocurrents between the full-length and the N-terminally truncated variant (13–428). **a** Fluorescence image of the N-terminally truncated variant (13–428) expressed in an ND7/23 cell. Scale bar: 20 μm. **b** Representative photocurrent traces of the full-length protein and the N-terminally truncated variant (13–428) recorded with standard pipette and extracellular solutions.

**Figure S7.**
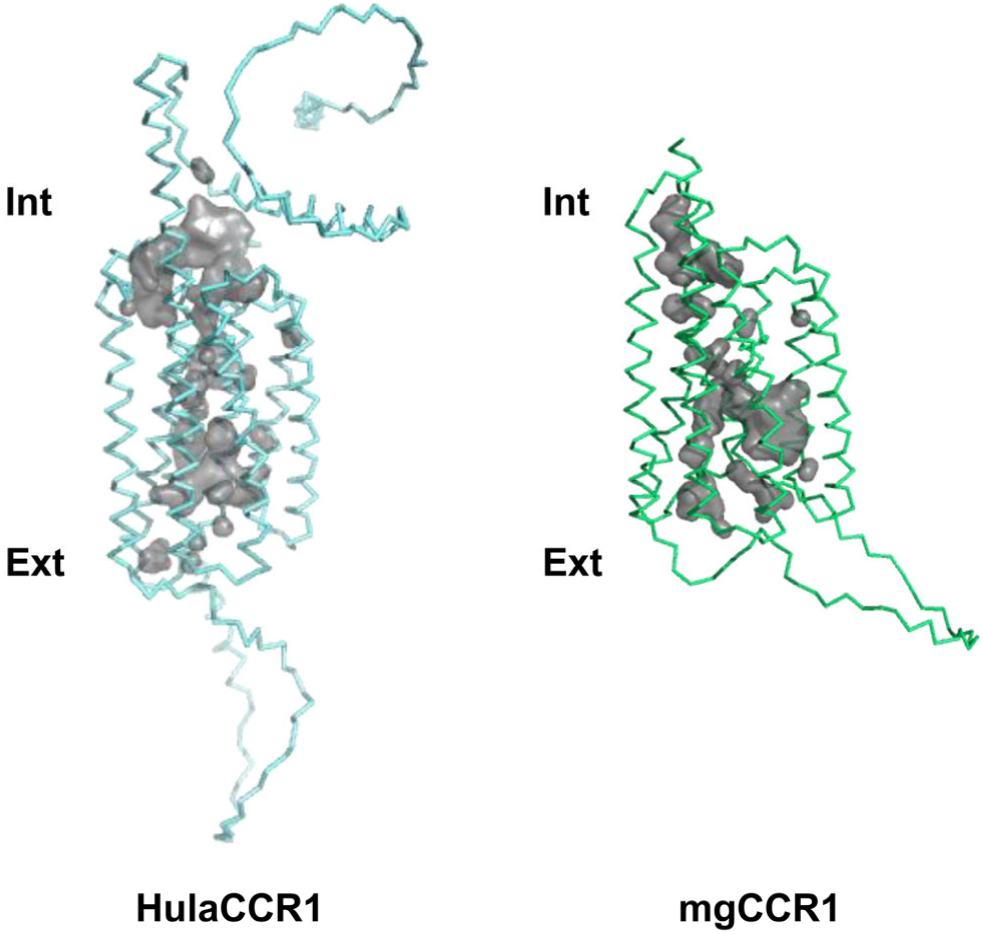
Internal cavities based on AlphaFold2 predicted structures of HulaCCR1 and mgCCR1. Calculated cavities by PyMOL software (version 2.6.0a0) are shown in gray.

**Table S1.**
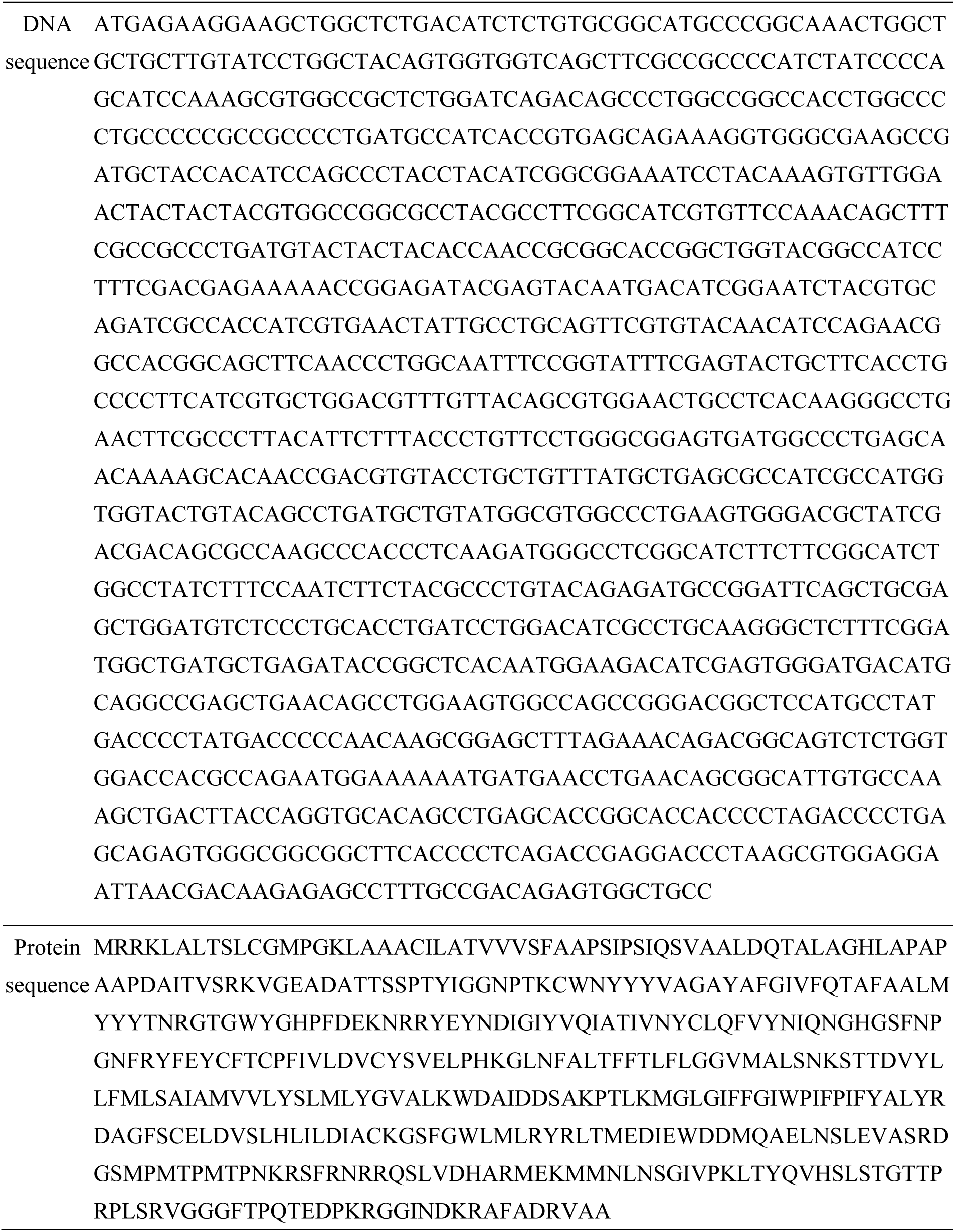
DNA and protein sequences of HulaCCR1.

**Table S2.**
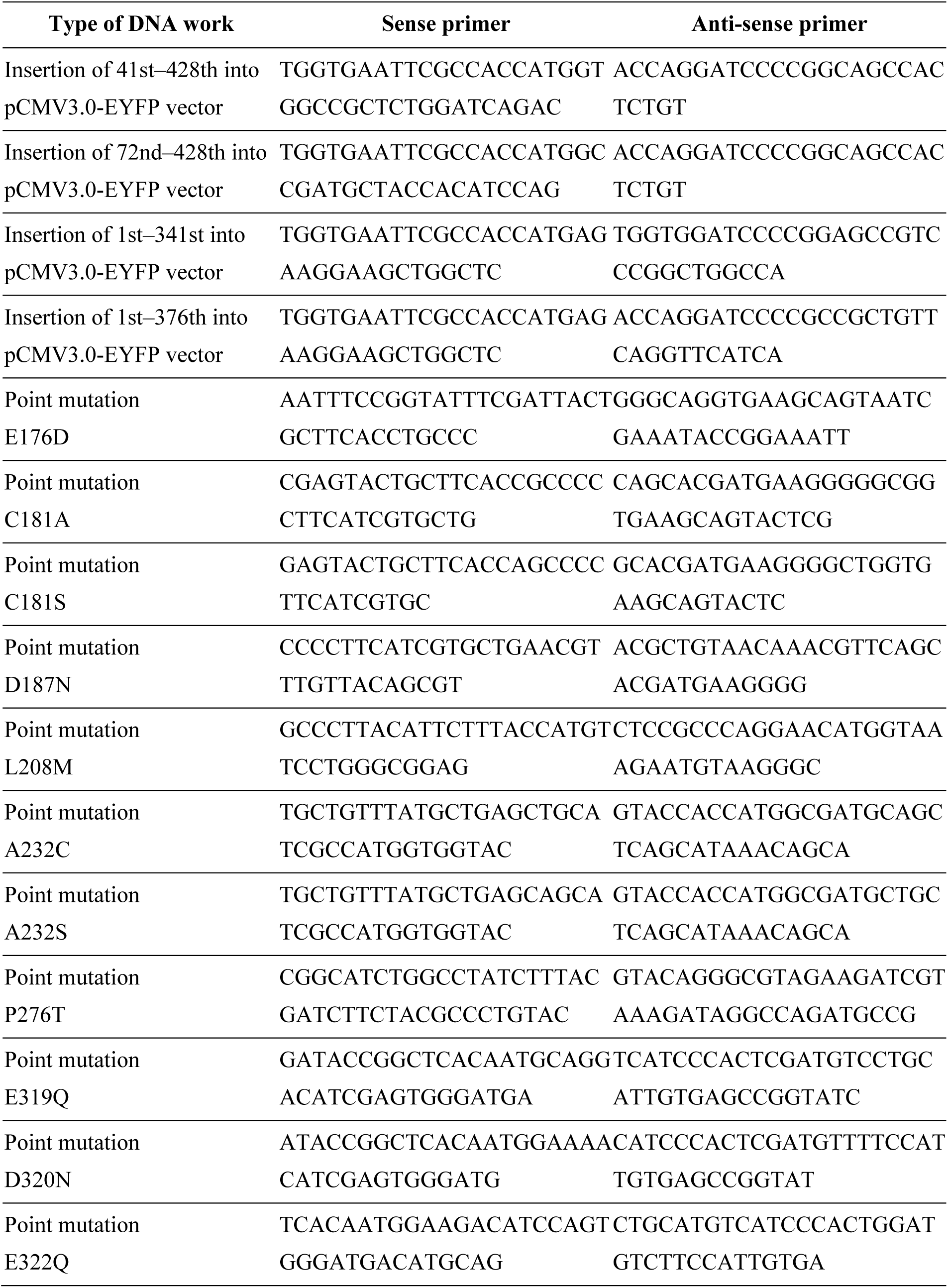
List of primer sequences used in DNA work.

**Table S3.**
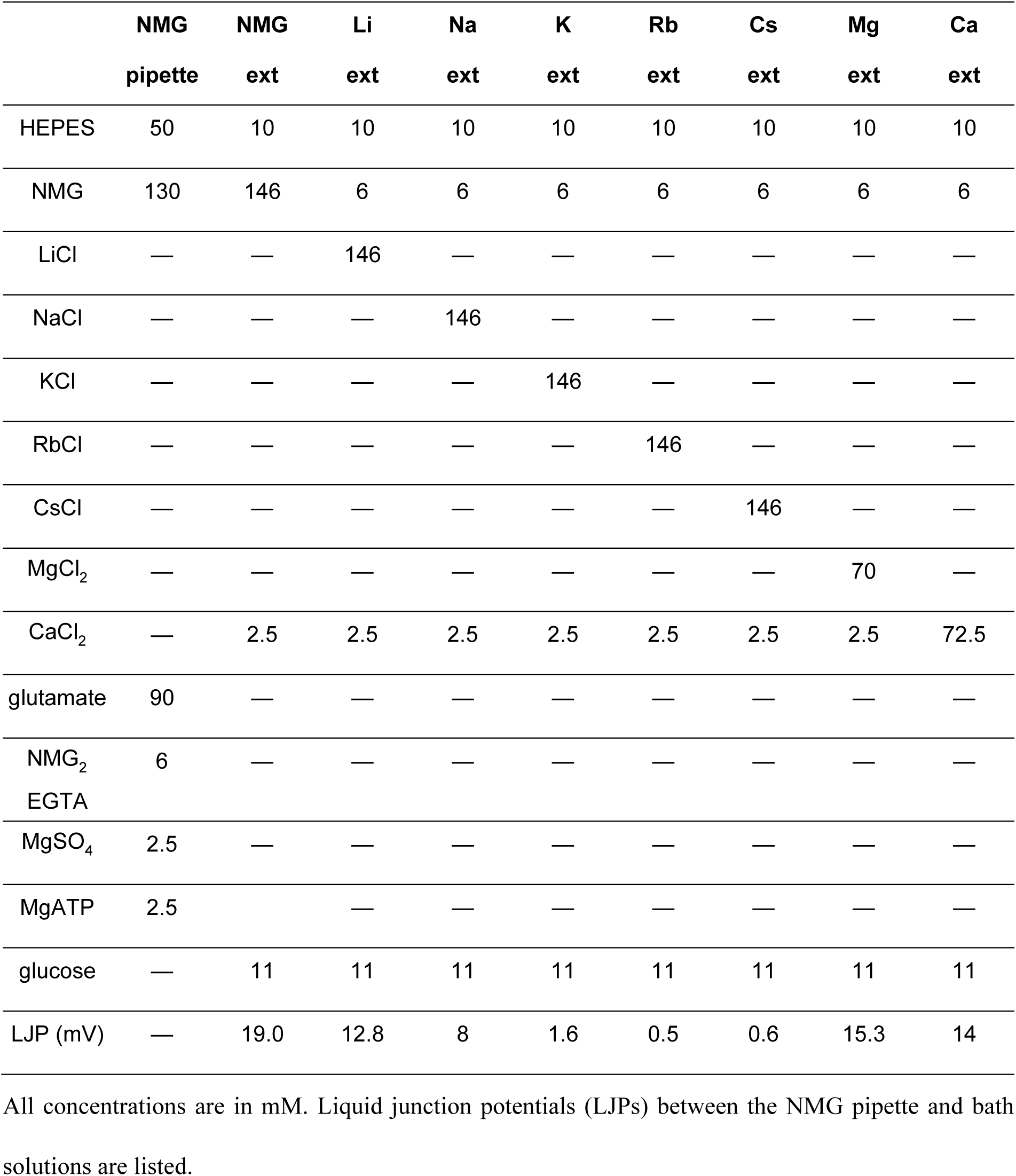
Composition of extracellular solutions used for ion-selectivity measurements.

## References

[1] A. Rozenberg, K. Inoue, H. Kandori, O. Béjà, Microbial rhodopsins: The last two decades, Annu. Rev. Microbiol. 75 (2021) 427–447.

[2] O.P. Ernst, D.T. Lodowski, M. Elstner, P. Hegemann, L.S. Brown, H. Kandori, Microbial and animal rhodopsins: structures, functions, and molecular mechanisms, Chem. Rev. 114 (2014) 126–163.

[3] T. Nagata, K. Inoue, Rhodopsins at a glance, J. Cell Sci. 134 (2021) jcs258989.

[4] K. Deisseroth, P. Hegemann, The form and function of channelrhodopsin, Science 357 (2017) eaan5544.

[5] V. Emiliani, E. Entcheva, R. Hedrich, P. Hegemann, K.R. Konrad, C. Lüscher, M. Mahn, Z.-H. Pan, R.R. Sims, J. Vierock, O. Yizhar, Optogenetics for light control of biological systems, Nat. Rev. Methods Primers 2 (2022) 55.

[6] O.A. Sineshchekov, K.-H. Jung, J.L. Spudich, Two rhodopsins mediate phototaxis to low– and high-intensity light in *Chlamydomonas reinhardtii*, Proc. Natl. Acad. Sci. U. S. A. 99 (2002) 8689–8694.

[7] G. Nagel, D. Ollig, M. Fuhrmann, S. Kateriya, A.M. Musti, E. Bamberg, P. Hegemann, Channelrhodopsin-1: a light-gated proton channel in green algae, Science 296 (2002) 2395– 2398.

[8] G. Nagel, T. Szellas, W. Huhn, S. Kateriya, N. Adeishvili, P. Berthold, D. Ollig, P. Hegemann, E. Bamberg, Channelrhodopsin-2, a directly light-gated cation-selective membrane channel, Proc. Natl. Acad. Sci. U. S. A. 100 (2003) 13940–13945.

[9] E.G. Govorunova, O.A. Sineshchekov, R. Janz, X. Liu, J.L. Spudich, Natural light-gated anion channels: A family of microbial rhodopsins for advanced optogenetics, Science 349 (2015) 647–650.

[10] E.G. Govorunova, O.A. Sineshchekov, J.L. Spudich, Structurally distinct cation channelrhodopsins from cryptophyte algae, Biophys. J. 110 (2016) 2302–2304.

[11] H. Hagio, W. Koyama, S. Hosaka, A.D. Song, J. Narantsatsral, K. Matsuda, T. Shimizu, S. Hososhima, S.P. Tsunoda, H. Kandori, M. Hibi, Optogenetic manipulation of neuronal and cardiomyocyte functions in zebrafish using microbial rhodopsins and adenylyl cyclases, eLife 12 (2023) e83975.

[12] M. Mahn, L. Gibor, P. Patil, K. Cohen-Kashi Malina, S. Oring, Y. Printz, R. Levy, I. Lampl, O. Yizhar, High-efficiency optogenetic silencing with soma-targeted anion-conducting channelrhodopsins, Nat. Commun. 9 (2018) 4125.

[13] S. Huang, M. Ding, M.R.G. Roelfsema, I. Dreyer, S. Scherzer, K.A.S. Al-Rasheid, S. Gao, G. Nagel, R. Hedrich, K.R. Konrad, Optogenetic control of the guard cell membrane potential and stomatal movement by the light-gated anion channel *Gt*ACR1, Sci. Adv. 7 (2021) eabg4619.

[14] Y. Zhou, M. Ding, S. Gao, J. Yu-Strzelczyk, M. Krischke, X. Duan, J. Leide, M. Riederer, M.J. Mueller, R. Hedrich, K.R. Konrad, G. Nagel, Optogenetic control of plant growth by a microbial rhodopsin, Nat. Plants 7 (2021) 144–151.

[15] H. Kandori, Ion-pumping microbial rhodopsins, Front. Mol. Biosci. 2 (2015) 52.

[16] Y. Yamauchi, M. Konno, S. Ito, S.P. Tsunoda, K. Inoue, H. Kandori, Molecular properties of a DTD channelrhodopsin from *Guillardia theta*, Biophys Physicobiol. 14 (2017) 57–66.

[17] J.H. Marshel, Y.S. Kim, T.A. Machado, S. Quirin, B. Benson, J. Kadmon, C. Raja, A. Chibukhchyan, C. Ramakrishnan, M. Inoue, J.C. Shane, D.J. McKnight, S. Yoshizawa, H.E. Kato, S. Ganguli, K. Deisseroth, Cortical layer-specific critical dynamics triggering perception, Science 365 (2019) eaaw5202.

[18] R. Chen, F. Gore, Q.-A. Nguyen, C. Ramakrishnan, S. Patel, S.H. Kim, M. Raffiee, Y.S. Kim, B. Hsueh, E. Krook-Magnusson, I. Soltesz, K. Deisseroth, Deep brain optogenetics without intracranial surgery, Nat. Biotechnol. 39 (2021) 161–164.

[19] H.E. Kato, Structure–Function Relationship of Channelrhodopsins, in: H. Yawo, H. Kandori, A. Koizumi, R. Kageyama (Eds.), Optogenetics: Light-sensing proteins and their applications in neuroscience and beyond, Springer Singapore, Singapore, 2021: pp. 35–53.

[20] E.G. Govorunova, Y. Gou, O.A. Sineshchekov, H. Li, X. Lu, Y. Wang, L.S. Brown, F. St-Pierre, M. Xue, J.L. Spudich, Kalium channelrhodopsins are natural light-gated potassium channels that mediate optogenetic inhibition, Nat. Neurosci. 25 (2022) 967–974.

[21] J. Vierock, E. Shiewer, C. Grimm, A. Rozenberg, I.-W. Chen, L. Tillert, A.G. Castro Scalise, M. Casini, S. Augustin, D. Tanese, B.C. Forget, R. Peyronnet, F. Schneider-Warme, V. Emiliani, O. Béjà, P. Hegemann, WiChR, a highly potassium-selective channelrhodopsin for low-light one– and two-photon inhibition of excitable cells, Sci. Adv. 8 (2022) eadd7729.

[22] E.G. Govorunova, O.A. Sineshchekov, L.S. Brown, A.-N. Bondar, J.L. Spudich, Structural foundations of potassium selectivity in channelrhodopsins, MBio 13 (2022) e0303922.

[23] M. Mirdita, K. Schütze, Y. Moriwaki, L. Heo, S. Ovchinnikov, M. Steinegger, ColabFold: making protein folding accessible to all, Nat. Methods 19 (2022) 679–682.

[24] J.G. Wideman, A. Monier, R. Rodríguez-Martínez, G. Leonard, E. Cook, C. Poirier, F. Maguire, D.S. Milner, N.A.T. Irwin, K. Moore, A.E. Santoro, P.J. Keeling, A.Z. Worden, T.A. Richards, Unexpected mitochondrial genome diversity revealed by targeted single-cell genomics of heterotrophic flagellated protists, Nat. Microbiol. 5 (2020) 154–165.

[25] N. Okamoto, I. Inouye, The katablepharids are a distant sister group of the Cryptophyta: A proposal for Katablepharidophyta divisio nova/ Kathablepharida phylum novum based on SSU rDNA and beta-tubulin phylogeny, Protist 156 (2005) 163–179.

[26] N. Okamoto, C. Chantangsi, A. Horák, B.S. Leander, P.J. Keeling, Molecular phylogeny and description of the novel katablepharid *Roombia truncata* gen. et sp. nov., and establishment of the Hacrobia taxon nov, PLoS One 4 (2009) e7080.

[27] J. Slapeta, P. López-García, D. Moreira, Present status of the molecular ecology of kathablepharids, Protist 157 (2006) 7–11.

[28] J. Oppermann, A. Rozenberg, T. Fabrin, C. GonzalezCabrera, O. Béjà, M. Prigge, P. Hegemann, Robust optogenetic inhibition with red-light-sensitive anion-conducting channelrhodopsins, eLife (2023). 10.7554/elife.90100.1.

[29] A. Rozenberg, A catalog of natural channelrhodopsins. Zenodo. 10.5281/zenodo.11306563

[30] K. Inoue, M. Karasuyama, R. Nakamura, M. Konno, D. Yamada, K. Mannen, T. Nagata, Y. Inatsu, H. Yawo, K. Yura, O. Béjà, H. Kandori, I. Takeuchi, Exploration of natural red-shifted rhodopsins using a machine learning-based Bayesian experimental design, Commun. Biol. 4 (2021) 362.

[31] C.A. Lewis, Ion-concentration dependence of the reversal potential and the single channel conductance of ion channels at the frog neuromuscular junction, J. Physiol. 286 (1979) 417– 445.

[32] S. Shigemura, S. Hososhima, H. Kandori, S.P. Tsunoda, Ion channel properties of a cation channelrhodopsin, Gt_CCR4, Appl. Sci. 9 (2019) 3440.

[33] K.E. Kishi, Y.S. Kim, M. Fukuda, M. Inoue, T. Kusakizako, P.Y. Wang, C. Ramakrishnan, E.F.X. Byrne, E. Thadhani, J.M. Paggi, T.E. Matsui, K. Yamashita, T. Nagata, M. Konno, S. Quirin, M. Lo, T. Benster, T. Uemura, K. Liu, M. Shibata, N. Nomura, S. Iwata, O. Nureki, R.O. Dror, K. Inoue, K. Deisseroth, H.E. Kato, Structural basis for channel conduction in the pump-like channelrhodopsin ChRmine, Cell 185 (2022) 672–689.e23.

[34] M. Nack, I. Radu, M. Gossing, C. Bamann, E. Bamberg, G.F. von Mollard, J. Heberle, The DC gate in Channelrhodopsin-2: crucial hydrogen bonding interaction between C128 and D156, Photochem. Photobiol. Sci. 9 (2010) 194–198.

[35] A. Berndt, O. Yizhar, L.A. Gunaydin, P. Hegemann, K. Deisseroth, Bi-stable neural state switches, Nat. Neurosci. 12 (2009) 229–234.

[36] E.G. Govorunova, O.A. Sineshchekov, H. Li, Y. Wang, L.S. Brown, J.L. Spudich, RubyACRs, nonalgal anion channelrhodopsins with highly red-shifted absorption, Proc. Natl. Acad. Sci. U. S. A. 117 (2020) 22833–22840.

[37] S.A. Tennigkeit, R. Karapinar, T. Rudack, M.-A. Dreier, P. Althoff, D. Eickelbeck, T. Surdin, M. Grömmke, M.D. Mark, K. Spoida, M. Lübben, U. Höweler, S. Herlitze, K. Gerwert, Design of an ultrafast G protein switch based on a mouse melanopsin variant, Chembiochem 20 (2019) 1766–1771.

[38] A.A. Shtyrov, D.M. Nikolaev, V.N. Mironov, A.V. Vasin, M.S. Panov, Y.S. Tveryanovich, M.N. Ryazantsev, Simple models to study spectral properties of microbial and animal rhodopsins: Evaluation of the electrostatic effect of charged and polar residues on the first absorption band maxima, Int. J. Mol. Sci. 22 (2021) 3029.

[39] K. Stehfest, E. Ritter, A. Berndt, F. Bartl, P. Hegemann, The branched photocycle of the slow-cycling channelrhodopsin-2 mutant C128T, J. Mol. Biol. 398 (2010) 690–702.

[40] O.A. Sineshchekov, E.G. Govorunova, H. Li, J.L. Spudich, Bacteriorhodopsin-like channelrhodopsins: Alternative mechanism for control of cation conductance, Proc. Natl. Acad. Sci. U. S. A. 114 (2017) E9512–E9519.

[41] S. Tajima, Y.S. Kim, M. Fukuda, Y. Jo, P.Y. Wang, J.M. Paggi, M. Inoue, E.F.X. Byrne, K.E. Kishi, S. Nakamura, C. Ramakrishnan, S. Takaramoto, T. Nagata, M. Konno, M. Sugiura, K. Katayama, T.E. Matsui, K. Yamashita, S. Kim, H. Ikeda, J. Kim, H. Kandori, R.O. Dror, K. Inoue, K. Deisseroth, H.E. Kato, Structural basis for ion selectivity in potassium-selective channelrhodopsins, Cell 186 (2023) 4325–4344.

[42] L.S. Brown, R. Needleman, J.K. Lanyi, Origins of deuterium kinetic isotope effects on the proton transfers of the bacteriorhodopsin photocycle, Biochemistry 39 (2000) 938–945.

[43] K. Inoue, S. Tahara, Y. Kato, S. Takeuchi, T. Tahara, H. Kandori, Spectroscopic study of proton-transfer mechanism of inward proton-pump rhodopsin, *Parvularcula oceani* Xenorhodopsin, J. Phys. Chem. B 122 (2018) 6453–6461.

[44] K. Shibata, K. Oda, T. Nishizawa, Y. Hazama, R. Ono, S. Takaramoto, R. Bagherzadeh, H. Yawo, O. Nureki, K. Inoue, H. Akiyama, Twisting and protonation of retinal chromophore regulate channel gating of channelrhodopsin C1C2, J. Am. Chem. Soc. 145 (2023) 10779– 10789.

[45] F. Teufel, J.J. Almagro Armenteros, A.R. Johansen, M.H. Gíslason, S.I. Pihl, K.D. Tsirigos, O. Winther, S. Brunak, G. von Heijne, H. Nielsen, SignalP 6.0 predicts all five types of signal peptides using protein language models, Nat. Biotechnol. 40 (2022) 1023–1025.

[46] K.D. Tsirigos, C. Peters, N. Shu, L. Käll, A. Elofsson, The TOPCONS web server for consensus prediction of membrane protein topology and signal peptides, Nucleic Acids Res. 43 (2015) W401–7.

[47] J. Mähler, I. Persson, A study of the hydration of the alkali metal ions in aqueous solution, Inorg. Chem. 51 (2012) 425–438.

[48] Y.S. Kim, H.E. Kato, K. Yamashita, S. Ito, K. Inoue, C. Ramakrishnan, L.E. Fenno, K.E. Evans, J.M. Paggi, R.O. Dror, H. Kandori, B.K. Kobilka, K. Deisseroth, Crystal structure of the natural anion-conducting channelrhodopsin *Gt*ACR1, Nature 561 (2018) 343–348.

[49] A. Rozenberg, J. Oppermann, J. Wietek, R.G. Fernandez Lahore, R.-A. Sandaa, G. Bratbak, P. Hegemann, O. Béjà, Lateral gene transfer of anion-conducting channelrhodopsins between green algae and giant viruses, Curr. Biol. 30 (2020) 4910–4920.e5.

[50] A. Kianianmomeni, K. Stehfest, G. Nematollahi, P. Hegemann, A. Hallmann, Channelrhodopsins of *Volvox carteri* are photochromic proteins that are specifically expressed in somatic cells under control of light, temperature, and the sex inducer, Plant Physiol. 151 (2009) 347–366.

[51] R. Tashiro, K. Sushmita, S. Hososhima, S. Sharma, S. Kateriya, H. Kandori, S.P. Tsunoda, Specific residues in the cytoplasmic domain modulate photocurrent kinetics of channelrhodopsin from *Klebsormidium nitens*, Commun. Biol. 4 (2021) 235.

[52] F. Krueger, F. James, P. Ewels, E Afyounian, M. Weinstein, B. Schuster-Boeckler, G. Hulselmans, FelixKrueger/TrimGalore: v0.6.10 – add default decompression path. Zenodo. 10.5281/zenodo.7598955

[53] M. Martin, Cutadapt removes adapter sequences from high-throughput sequencing reads, EMBnet.Journal 17 (2011) 10–12.

[54] D. Li, C.-M. Liu, R. Luo, K. Sadakane, T.-W. Lam, MEGAHIT: an ultra-fast single-node solution for large and complex metagenomics assembly via succinct de Bruijn graph, Bioinformatics 31 (2015) 1674–1676.

[55] W. Li, A. Godzik, Cd-hit: a fast program for clustering and comparing large sets of protein or nucleotide sequences, Bioinformatics 22 (2006) 1658–1659.

[56] L.S. Johnson, S.R. Eddy, E. Portugaly, Hidden Markov model speed heuristic and iterative HMM search procedure, BMC Bioinformatics 11 (2010) 431.

[57] I.-M.A. Chen, K. Chu, K. Palaniappan, A. Ratner, J. Huang, M. Huntemann, P. Hajek, S.J. Ritter, C. Webb, D. Wu, N.J. Varghese, T.B.K. Reddy, S. Mukherjee, G. Ovchinnikova, M. Nolan, R. Seshadri, S. Roux, A. Visel, T. Woyke, E.A. Eloe-Fadrosh, N.C. Kyrpides, N.N. Ivanova, The IMG/M data management and analysis system v.7: content updates and new features, Nucleic Acids Res. 51 (2023) D723–D732.

[58] C. Camacho, G. Coulouris, V. Avagyan, N. Ma, J. Papadopoulos, K. Bealer, T.L. Madden, BLAST+: architecture and applications, BMC Bioinformatics 10 (2009) 421.

[59] K. Katoh, D.M. Standley, MAFFT multiple sequence alignment software version 7: improvements in performance and usability, Mol. Biol. Evol. 30 (2013) 772–780.

[60] A. Stamatakis, RAxML version 8: a tool for phylogenetic analysis and post-analysis of large phylogenies, Bioinformatics 30 (2014) 1312–1313.

[61] F.A. Matsen, R.B. Kodner, E.V. Armbrust, pplacer: linear time maximum-likelihood and Bayesian phylogenetic placement of sequences onto a fixed reference tree, BMC Bioinformatics 11 (2010) 538.

[62] L. Czech, P. Barbera, A. Stamatakis, Genesis and Gappa: processing, analyzing and visualizing phylogenetic (placement) data, Bioinformatics 36 (2020) 3263–3265.

[63] L. Fu, B. Niu, Z. Zhu, S. Wu, W. Li, CD-HIT: accelerated for clustering the next-generation sequencing data, Bioinformatics 28 (2012) 3150–3152.

[64] D.J. Richter, C. Berney, J.F.H. Strassert, Y.-P. Poh, E.K. Herman, S.A. Muñoz-Gómez, J.G. Wideman, F. Burki, C. de Vargas, EukProt: A database of genome-scale predicted proteins across the diversity of eukaryotes, Peer Community J. 2 (2022) e56.

[65] T.O. Delmont, M. Gaia, D.D. Hinsinger, P. Fremont, C. Vanni, A.F. Guerra, A. Murat Eren, A. Kourlaiev, L. d’Agata, Q. Clayssen, E. Villar, K. Labadie, C. Cruaud, J. Poulain, C. Da Silva, M. Wessner, B. Noel, J.-M. Aury, T.O. Coordinators, C. de Vargas, C. Bowler, E. Karsenti, E. Pelletier, P. Wincker, O. Jaillon, Functional repertoire convergence of distantly related eukaryotic plankton lineages revealed by genome-resolved metagenomics, BioRxiv (2021) 2020.10.15.341214. 10.1101/2020.10.15.341214.

[66] S. Capella-Gutiérrez, J.M. Silla-Martínez, T. Gabaldón, trimAl: a tool for automated alignment trimming in large-scale phylogenetic analyses, Bioinformatics 25 (2009) 1972– 1973.

[67] R.C. Edgar, SINTAX: a simple non-Bayesian taxonomy classifier for 16S and ITS sequences, BioRxiv (2016) 074161. 10.1101/074161.

[68] L. Guillou, D. Bachar, S. Audic, D. Bass, C. Berney, L. Bittner, C. Boutte, G. Burgaud, C. de Vargas, J. Decelle, J. Del Campo, J.R. Dolan, M. Dunthorn, B. Edvardsen, M. Holzmann, W.H.C.F. Kooistra, E. Lara, N. Le Bescot, R. Logares, F. Mahé, R. Massana, M. Montresor, R. Morard, F. Not, J. Pawlowski, I. Probert, A.-L. Sauvadet, R. Siano, T. Stoeck, D. Vaulot, P. Zimmermann, R. Christen, The Protist Ribosomal Reference database (PR2): a catalog of unicellular eukaryote small sub-unit rRNA sequences with curated taxonomy, Nucleic Acids Res. 41 (2013) D597–604.

[69] V. Gradinaru, F. Zhang, C. Ramakrishnan, J. Mattis, R. Prakash, I. Diester, I. Goshen, K.R. Thompson, K. Deisseroth, Molecular and cellular approaches for diversifying and extending optogenetics, Cell 141 (2010) 154–165.

[70] N. Morimoto, T. Nagata, K. Inoue, Reversible photoreaction of a retinal photoisomerase, retinal G-protein-coupled receptor RGR, Biochemistry 62 (2023) 1429–1432.

[71] J.F. Fay, D.L. Farrens, Chapter fifteen – Purification of functional CB1 and analysis by site-directed fluorescence labeling methods, in: P.H. Reggio (Ed.), Methods in Enzymology, Academic Press, 2017: pp. 343–370.

## References

[1] J.D. Thompson, D.G. Higgins, T.J. Gibson, CLUSTAL W: improving the sensitivity of progressive multiple sequence alignment through sequence weighting, position-specific gap penalties and weight matrix choice, Nucleic Acids Res. 22 (1994) 4673–4680.

[2] X. Robert, P. Gouet, Deciphering key features in protein structures with the new ENDscri pt server, Nucleic Acids Res. 42 (2014) W320–4.

